# ImmTACs overcome cytotoxic T cell suppression

**DOI:** 10.64898/2026.03.20.713194

**Authors:** Lan Huynh, Abdullah Aljohani, Amal Alsubaiti, Tressan Grant, Alexandra Chapman, Gwilym Phillips, Jonathan Chamberlain, Alice Hayward-Wills, Ute Jungwirth, Mariolina Salio, Christopher J. Holland, Christoph Wülfing

**Author notes:** equal contribution (Lan Huynh, Abdullah Aljohani, Amal Alsubaiti).

## Abstract

Immune mobilizing monoclonal TCR against cancer (ImmTAC) are cancer therapeutics that activate T cells through recognition of a tumor-associated antigenic MHC/peptide complex. A first-in-class ImmTAC, Tebentafusp, is approved for the treatment of metastatic uveal melanoma. While clinical efficacy is thus established, the cellular mechanisms underpinning ImmTAC action are not fully resolved. Using a recently established experimental strategy to generate suppressed human primary cytotoxic T lymphocytes (CTL), we have investigated an ImmTAC that recognizes a peptide derived from the tumor associated antigen NY-ESO-1 in comparison to direct engagement of a TCR recognizing the same MHC/peptide complex. In response to endogenous antigen presentation, ImmTACs could elicit tumor cell cytolysis by suppressed CTL, but not IFNγ secretion, in a manner dependent on the engager affinity for CD3ε. ImmTACs enhanced the efficient execution of subcellular CTL polarization steps required for effective cytolysis and could trigger calcium signaling. These data establish that ImmTACs activate CTL similarly to direct engagement of a TCR by MHC/peptide and are likely to retain this capability under suppressive conditions such as in the tumor microenvironment.

## Introduction

Cancer commonly triggers an anti-tumor immune response with the potential to cure the disease. Key immune effectors are cytotoxic T lymphocytes (CTL) that can directly kill tumor target cells [1]. To do so, the T cell receptor (TCR) of a CTL needs to recognize a peptide derived from a tumor-associated or neoantigen as presented by Major Histocompatibility (MHC) molecules on the tumor cell surface. Common challenges to an effective T cell-mediated anti-tumor immune response are insufficient numbers of tumor-reactive T cells and a suppressive tumor microenvironment (TME) that impairs the function of tumor-infiltrating T cells. Multiple therapeutic strategies aim to increase the number of T cells that can be activated in the TME. These include adoptive cell therapies with autologous in vitro expanded tumor-infiltrating lymphocytes and autologous T cells virally engineered to express a tumor-reactive TCR or a chimeric antigen receptor [2, 3]. Alternatively, synthetic T cell engagers aim to activate all T cells present in the TME [4]. As a substantial fraction of T cells in the TME are bystander T cells [5, 6], i.e. don’t express a tumor antigen-reactive TCR, this strategy can substantially increase the number of activated T cells. Such engagers are commonly bivalent proteins that bind to an antigen expressed on the tumor cell surface and a subunit of the TCR.

The initial design of bispecific T cell engagers connects a tumor-specific cell surface protein recognized with an antibody fragment to the CD3ε subunit of the TCR [4, 7]. However, with only 10% of a cell’s proteome accessible on the cell surface, the number of proteins that can be recognized by such engagers can be limiting. To access the intracellular proteome for tumor cell recognition, Immune mobilizing monoclonal TCRs against cancer (ImmTAC) recognize MHC/peptide complexes on the tumor cell surface. ImmTACs are bispecific T cell engagers composed of the disulfide-linked extracellular domains of the alpha and beta chains of an affinity enhanced TCR recognizing a tumor associated antigenic peptide presented by MHC linked to an scFv fragment of an antibody against CD3ε [8]. Tebentafusp, an ImmTAC recognizing a peptide derived from gp100 as presented by HLA-A*0201, is clinically approved for the treatment of metastatic uveal melanoma [9, 10]. Patient data show enhanced expression of molecules associated with T cell activation in tumor biopsies [11]. This is consistent with in vitro data showing induction of tumor cell killing and T cell IFNγ secretion by ImmTACs [12]. However, questions about the mechanism of action of ImmTACs remain unresolved. It remains to be defined whether ImmTACs directly activate suppressed T cells present in the TME or they act indirectly, e.g. by enhancing T cell expansion in tumor-draining lymph nodes similar to immune checkpoint inhibitors [13]. Similarly, ImmTACs can activate CTL in vitro [14] but it is not clear whether this would extend to suppressed CTL, mimicking the CTL in the TME. Effective CTL-mediated target cell killing requires the sequential execution of a series of subcellular polarization steps to stabilize the CTL target cell couple [15–17]. Do ImmTACs regulate such subcellular polarization? Proximal T cell signaling, especially the elevation of the cytoplasmic calcium concentration, is associated with effective cytokine secretion in CTL [18]. Do ImmTACs regulate such signaling? Using three-dimensional tissue culture, we have recently established a strategy to generate suppressed CTL in vitro that closely resemble exhausted CTL in the TME [19]. We have employed this approach to address the above questions regarding the mechanism of action of ImmTAC molecules.

Synthetic design approaches have recently been applied to generate new specific binders to MHC/peptide complexes, including binders to the same MHC/peptide complex investigated here [20–22]. In their current iteration, these binders have affinities for the MHC/peptide complexes in the low nM range. They can be incorporated into synthetic TCR engagers analogous to an ImmTAC and have been shown to trigger effective T cell activation in this form [20]. Investigating the mechanism of action of ImmTACs thus should have broad applicability [23]. We use an ImmTAC recognizing the peptide 157-165 of the tumor-associated antigen NY-ESO-1 presented by HLA-A*0201 [23] as a model for this class of therapeutics and compare its activity to direct activation of CTL lentivirally transduced to express the 1G4 TCR that recognizes the same peptide/MHC complex [24]. As effectors, we use in vitro suppressed CTL generated following activation of primary human CTL with tumor cell spheroids [19]. As targets, we use the melanoma cell lines Mel624 and A375, both of which express low amounts of NY-ESO-1 [19]. We show that under suppressive conditions ImmTACs are effective at eliciting cytolysis but not IFNγ secretion, and this process is fine-tuned by anti-CD3ε affinity [12]. Finally, using live cell imaging of the interaction of suppressed CTL with tumor target cells we show that ImmTACs behave similarly to direct TCR engagement by MHC/peptide in enhancing the subcellar polarization of CTL and leading to elevation of the intracellular calcium concentration. Together, our data suggest that ImmTACs are effective reagents to trigger cytolysis even under suppressive conditions with a mechanism that closely resembles direct TCR engagement by MHC/peptide.

## Results

### ImmTACs effectively trigger cytolysis in response to endogenous amounts of antigen

As reference data for an investigation of ImmTACs in a suppressive environment, we first determined the ability of ImmTACs to activate CTL as a function of antigen dose, ImmTAC dose and ImmTAC affinity for CD3ε. We used ImmTAC molecules that recognize the peptide 157-165 of the tumor-associated antigen NY-ESO-1 presented by HLA-A*0201 [23]. We used four ImmTACs differing in binding to CD3ε with affinity increasing from E8<E0=E28<E42 and with faster on and off rates of E28 than E0 [12](Fig. 1A). We used human in vitro primed CTL as previously characterized [19], per se (‘CD8^+^ CTL’) or lentivirally transduced to express the 1G4 TCR that recognizes the same NY-ESO-1_157-165_/HLA-A*0201 complex (‘1G4 CTL’) [24] (Fig. 1A) to allow a comparison with CTL activation by direct engagement of the TCR by MHC/peptide. As tumor target cells, we predominantly used Mel624 melanoma cells, and in a few experiments A375 melanoma cells. Both cell lines express small amounts of NY-ESO-1 that allow for limiting activation of 1G4 CTL [19]. To generate saturating amounts of antigen, we have incubated the cell lines with 2 µg/ml of exogenous NY-ESO-1_157-165_ peptide (Fig. 1A). Mel624 cells can effectively be grown as spheroids (Fig. 1A) and incubation of such spheroids with 1G4 CTL induces suppression comparable to CTL exhaustion in tumors [19].

**Fig. 1.**
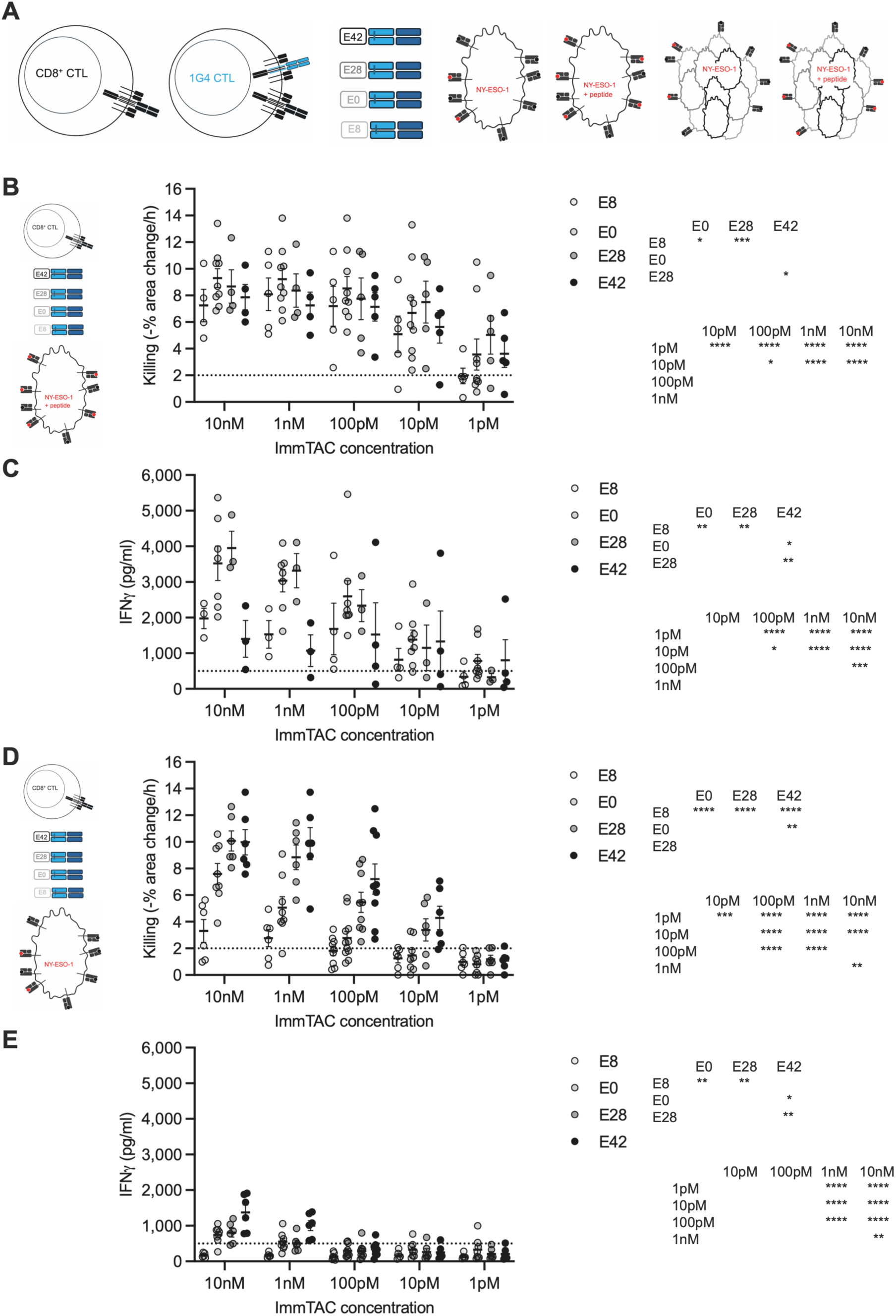
ImmTACs trigger CTL effector function in a dose and affinity-dependent fashion. **A** Graphical depiction of cell types and ImmTACs used, non-transduced CTL, CTL transduced to express the 1G4 TCR, ImmTACs with different anti-CD3ε scFv fragments, Mel624 or A375 melanoma cells with only endogenous NY-ESO-1 processing, Mel624 or A375 melanoma cells in the presence of 2 µg/ml NY-ESO-1_157-165_ peptide and the corresponding Mel 624 spheroids. Red dot indicates the NY-ESO-1_157-165_ peptide. **B** Killing of Mel624 or A375 tumor target cells (incubated with 2 µg/ml NY-ESO-1 peptide) by non-transduced CTL in the presence of the indicated concentration of the given ImmTAC version; data pooled for Mel624 and A375 tumor target cell; mean ± SEM. A broken line indicates the expected level of killing in the absence of antigen. 5-9 independent experiments. Statistical significance determined by paired Mixed-effects analysis and given as main effects of ImmTAC version and concentration in the two tables at the right. **C** IFNγ amounts in supernatants of Mel624 or A375 tumor target cells incubated with 2 µg/ml NY-ESO-1 peptide and interacting with non-transduced CTL in the presence of the indicated concentration of the given ImmTAC version; data pooled; mean ± SEM. A broken line indicates the expected level of IFNγ secretion in the absence of antigen. 3-8 independent experiments. Statistical significance determined by paired Mixed-effects analysis and given as in B. **D** Killing of Mel624 or A375 tumor target cells by non-transduced CTL in the presence of the indicated concentration of the given ImmTAC version; mean ± SEM. 9-11 independent experiments with not all ImmTAC concentrations present in all experiments. Statistical significance determined by Mixed-effects analysis and given as in B. **E** IFNγ amounts in supernatants of Mel624 or A375 tumor target cells interacting with non-transduced CTL in the presence of the indicated concentration of the given ImmTAC version; mean ± SEM. 7-10 independent experiments with not all ImmTAC concentrations present in all experiments. Statistical significance determined by paired Mixed-effects analysis and given as in B. * p<0.05, ** p<0.01, *** p<0.001, **** p<0.0001.

In response to target cells incubated with exogenous NY-ESO-1 peptide, all four ImmTACs triggered cytolysis (Fig. 1B) with only E8 being less effective, as determined in an imaging-based time-resolved overnight killing assay [16]. Cytolysis was saturated at ImmTAC concentrations of 10 pM or 100 pM. In patients, ImmTACs are given weekly leading to fluctuating in vivo concentrations [23]. In a phase 1 clinical trial of a comparable NY-ESO-1-directed ImmTAC as used here, ImmTAC plasma concentrations reach 50 pM at the time of administration [23] with uncertain uptake into tumors. We, therefore, use 100 pM in single concentration experiments. IFNγ secretion, as determined in the supernatants at the end of the killing assay, was less effectively triggered and showed greater differences between the different ImmTAC versions. IFNγ secretion was most efficiently triggered by the E0 and E28 ImmTACs, requiring between 100 pM and 1 nM for saturation (Fig. 1C). Less effective induction of IFNγ secretion in comparison to cytolysis is similarly found in the direct activation of 1G4 CTL by the same melanoma cells [19]. A bell-shaped dose response curve for receptor agonists as a function of affinity has also been previously described, explaining the results with E42 ImmTAC [12, 25].

CTL activation by ImmTACs in response to endogenous antigen presentation was substantially less effective. The E8 ImmTAC could no longer trigger effective cytolysis across the entire ImmTAC dose range (Fig. 1D). E28 and E42 were most effective but now required concentrations of between 100 pM and 1 nM for saturating cytolysis, an approximate 10-fold shift in the dose response curve compared to the cytolysis in the presence of saturating amounts of peptide (Fig. 1B). Such cytolysis was still antigen-dependent as shown with NY-ESO-1 knock down in Mel624 cells [19] (Fig. S1). Killing only occurred at a reduced level and at ImmTAC concentrations above 100 pM for E42 ImmTAC and at 10 nM for E28 and E0 molecules (Fig. S1A). In contrast to cytolysis, ImmTACs barely triggered IFNγ secretion in response to endogenous levels of NY-ESO-1. Amounts of IFNγ were substantially smaller across the entire dose range, and the minimal ImmTAC concentration to trigger any IFNγ secretion was 1 nM (Fig. 1E). The E8 ImmTAC did not induce any IFNγ secretion anymore (Fig. 1E). Comparable results were observed for IFNγ secretion in response to the NY-ESO-1 knock down Mel624 cells, with a 1 nM threshold for E42 and 10 nM for U28 (Fig. S1B).

Finally, we determined whether CD3ε engagement with ImmTACs synergizes with direct engagement of the α/β heterodimer of a TCR by MHC/peptide on the same T cell by comparing CD8^+^ CTL and 1G4 activation in response to 100 pM E28 ImmTAC. While killing by 1G4 CTL in response to E28 was slightly higher than that by CD8^+^ CTL, the difference did not reach statistical significance and is unlikely to be functionally relevant (Fig. S1C). IFNγ secretion was low with no substantial differences (Fig. S1D).

Together, these data establish that E28 and E42 ImmTACs at the most likely to be physiologically relevant concentration of 100 pM can trigger efficient cytolysis in response to limiting endogenous antigen presentation but without noticeable IFNγ secretion and irrespective of whether the CTL express a tumor-reactive TCR. Larger amounts of antigen enabled cytolytic responses by smaller amounts of ImmTAC, by ImmTACs with a weaker affinity for CD3ε and effective IFNγ secretion.

### The efficiency of CTL activation by ImmTACs is comparable to that triggered by direct TCR engagement by MHC/peptide

To gauge the efficacy of ImmTACs in comparison to physiological CTL activation through direct TCR engagement by MHC/peptide, we used 1G4 CTL (Fig. 2A). In response to Mel624 cells incubated with saturating amounts of exogenous NY-ESO-1 peptide, CTL activation with 100 pM ImmTACs triggered cytolysis that was comparable if not superior to that of 1G4 CTL (Fig. 2B). IFNγ secretion triggered by 100 pM E28 ImmTAC was greater than that of 1G4 CTL (Fig. 2C). In response to endogenous NY-ESO-1 antigen processing, 100 pM E0, E28 and E42 again triggered comparable if not greater cytolysis (Fig. 2D). There was no IFNγ secretion under these conditions (Fig. 2E). In the interpretation of these data, it is important to consider that the 1G4 CTL express the 1G4 TCR in addition to endogenous TCRs and as stabilized with only a disulfide bridge in the constant domains [26], as used in adoptive cell therapy. This leads to comparatively low 1G4 TCR expression and less effective CTL activation through the 1G4 CTL as should be expected in CTL only expressing the transgenic TCR [19]. With this caveat in mind, the data still establish that CTL activation by the E28 and E42 ImmTACs at the concentration most likely to be physiologically relevant of 100 pM is comparable to that of CTL activation by direct engagement of the TCR by MHC/peptide.

**Fig. 2.**
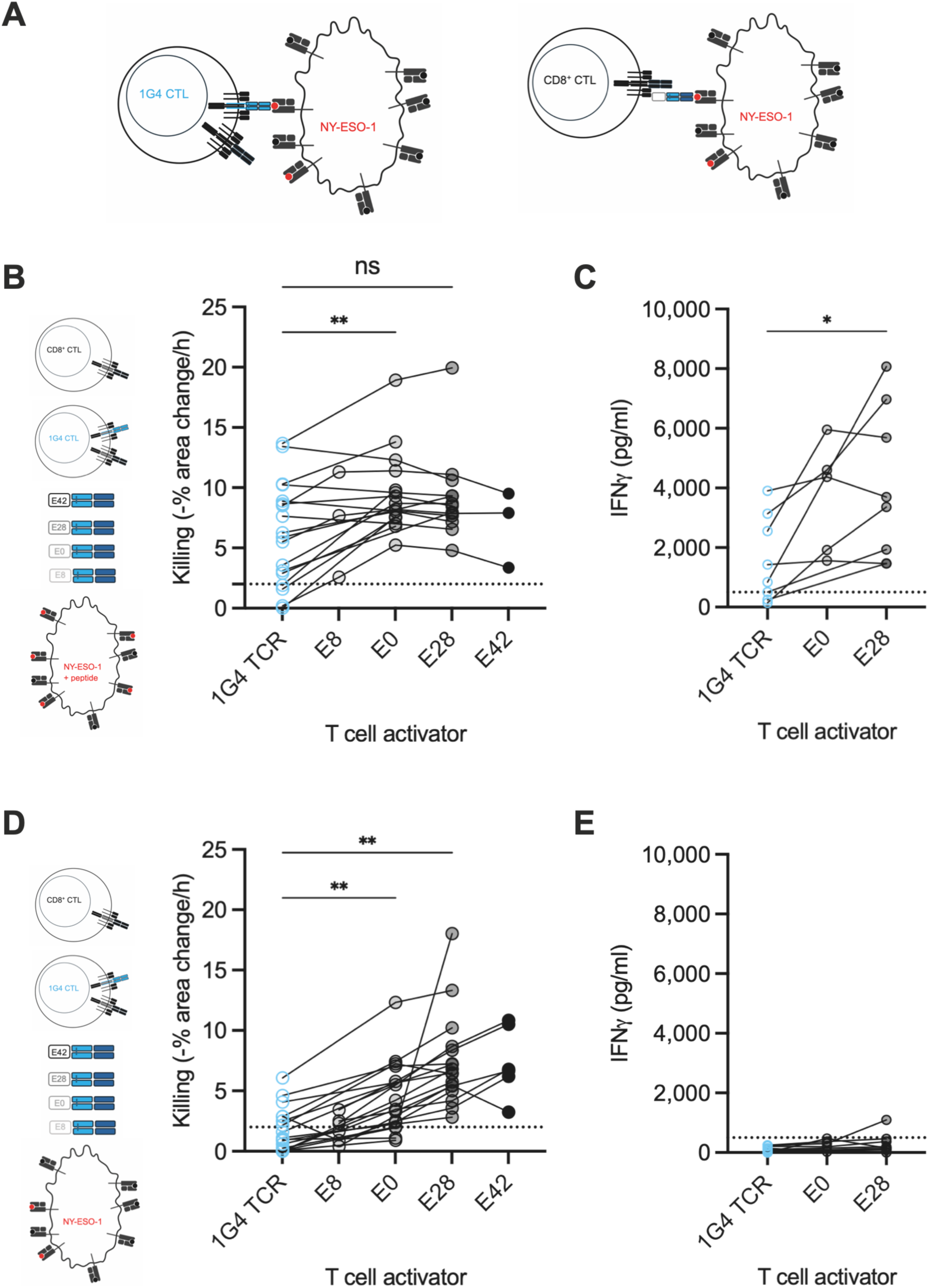
ImmTACs are more potent than T cell activation by direct engagement of the TCR by MHC/peptide. **A** Graphical depiction of the direct activation of 1G4 CTL by NY-ESO-1_157-165_/HLA-A*0201 versus the indirect activation of non-transduced CTL via ImmTACs recognizing the same NY-ESO-1_157-165_/HLA-A*0201 complex. **B** Killing of Mel624 or A375 tumor target cells incubated with 2 µg/ml NY-ESO-1 peptide by 1G4 CTL or non-transduced CTL in the absence or presence of 100 pM of the given ImmTAC version; data pooled; mean ± SEM. 3-18 independent experiments. Statistical significance determined by paired One-way ANOVA. **C** IFNγ amounts in supernatants of Mel624 or A375 tumor target cells incubated with 2 µg/ml NY-ESO-1 peptide interacting with 1G4 CTL or non-transduced CTL in the absence or presence of 100 pM of the given ImmTAC version; data pooled; mean ± SEM. 5-8 independent experiments. Statistical significance determined by paired One-way ANOVA. **D** Killing of Mel624 or A375 tumor target cells by 1G4 CTL or non-transduced CTL in the absence or presence of 100 pM of the given ImmTAC version; data pooled; mean ± SEM. 6-17 independent experiments. Statistical significance determined by paired One-way ANOVA. **E** IFNγ amounts in supernatants of Mel624 or A375 tumor target cells interacting with 1G4 CTL or non-transduced CTL in the absence or presence of 100 pM of the given ImmTAC version; data pooled; mean ± SEM. 5-9 independent experiments. Statistical significance determined by paired One-way ANOVA. * p<0.05, ** p<0.01.

### ImmTACs trigger tumor cell killing in a suppressive environment

In patients, ImmTACs must function in the suppressive tumor microenvironment. We use tumor cell spheroids to generate a suppressed CTL phenotype that is comparable to exhausted CTL in tumors [19]. Tumor cell killing is determined as the increase in dead spheroid volume in live spheroid imaging of Mel624 spheroids interacting with CTL [19](Fig. 3A). In response to Mel624 spheroids incubated with saturating amounts of exogenous NY-ESO-1 peptide, 100 pM E28 and less so E42 triggered substantial time-dependent tumor cell killing (Figs. 3B, S2A). This was accompanied by matching CTL infiltration into the spheroids (Fig. 3C) and a time-dependent increase in the depth of infiltration (Fig. S2B). In response to endogenous NY-ESO-1 antigen processing by the Mel624 cells, only 100 pM E42 ImmTAC could trigger cytolysis (Fig. 3D, S2C). Spheroid infiltration was reduced in response to all ImmTACs but still highest for E42 (Fig. 3E). Infiltration depth was largely unchanged (Fig. S2D). The extent of killing triggered by the ImmTAC E0 was comparable to killing triggered by direct TCR engagement with MHC/peptide in 1G4 CTL (Fig. S3). Together these data establish that ImmTACs at the concentration most likely to be physiologically relevant of 100 pM can trigger CTL infiltration into tumor cell spheroids leading to target cytolysis dependent on anti CD3 affinity, particularly in response to limiting endogenous amounts of antigen. CTL infiltration into spheroids and tumor cell killing were only partially related. Some infiltration occurred without triggering cytolysis. Going forward, we will focus on E28 ImmTAC molecules and occasionally include E0 to determine the role of different on/off rates.

**Fig. 3.**
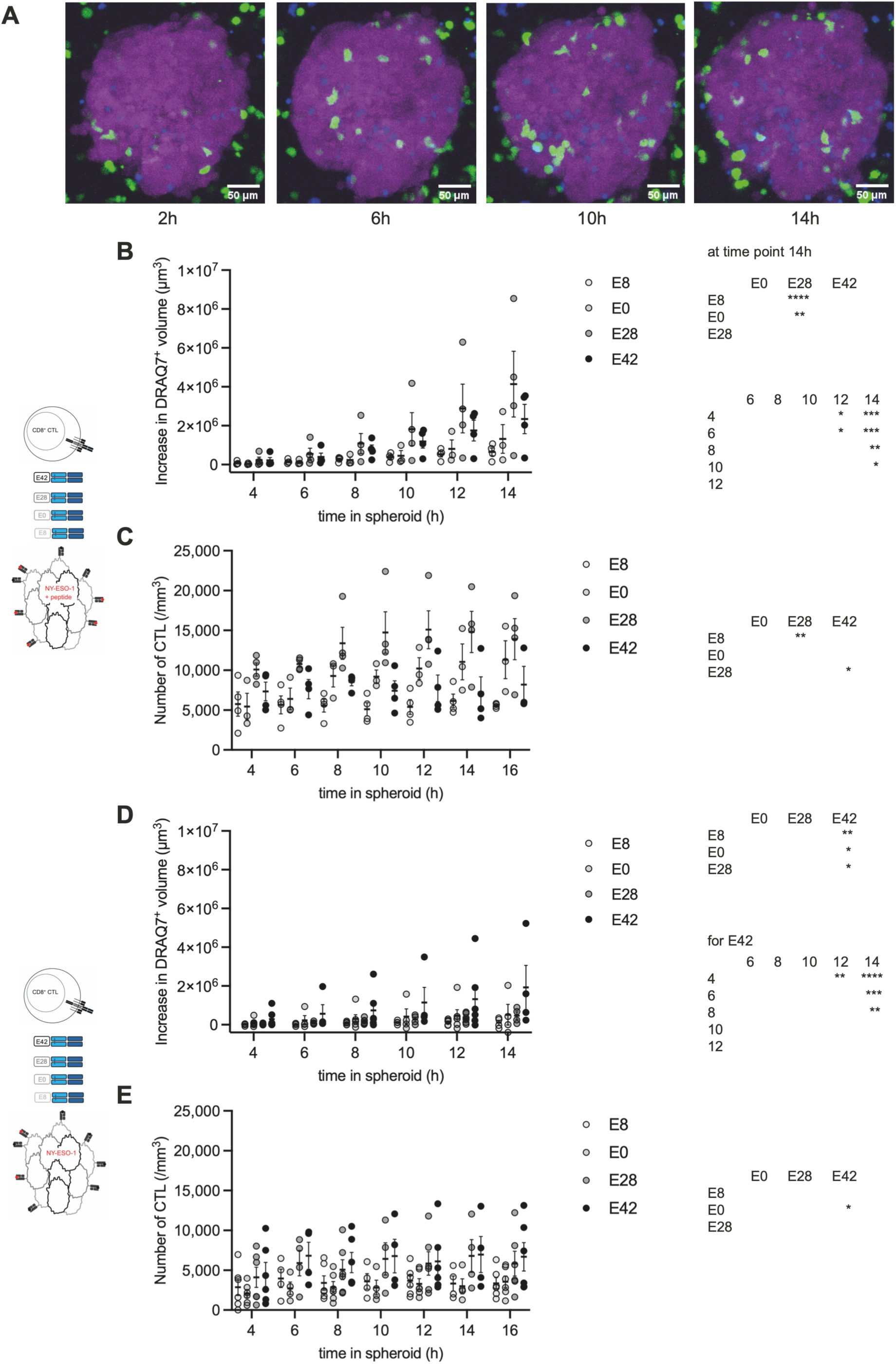
ImmTACs mediate target cell killing in tumor cell spheroids. **A** Representative spheroid killing data relative to the time of Matrigel embedding of Mel624 spheroids (purple) in the presence of exogenous NY-ESO-1 peptide with 100 pM E28, CD8^+^ CTL (green) and DRAQ7 staining (blue) to determine cell death. Maximum projection of 3D imaging data. Scale bar=50µm. **B, C** Non-transduced CTL cocultured with Mel624 spheroids incubated with 2 µg/ml NY-ESO-1 peptide. Each data point is an independent experiment (N=3, 4) with a total of 8-11 spheroids analyzed per condition. (B) Spheroid death, as measured by the increase in DRAQ7^+^ spheroid volume, is shown. Statistical significance determined by paired Mixed-effects analysis and given as differences between ImmTAC versions at 14 h and main effect of time in the tables at the right. Single spheroid data are given in Fig. S2A. (C) SIL densities are shown with the mean ± SEM for the same experiments as in B. Statistical significance determined by paired Mixed-effects analysis and given as main effect of ImmTAC version in the table at the right. **D, E** Non-transduced CTL cocultured with Mel624 spheroids for 12h with images acquired every 2 h or 4 h. Each data point is an independent experiment (N=6) with a total of 14-16 spheroids analyzed per condition. (D) Spheroid death, as measured by the increase in DRAQ7^+^ spheroid volume, is shown. Statistical significance determined by paired Mixed-effects analysis and given as differences between time points for E42 and main effect of ImmTAC version in the tables at the right. Single spheroid data are given in Fig. S2C. (E) SIL densities are shown with the mean ± SEM for the same experiments as in D. Statistical significance determined by paired Mixed-effects analysis and given as main effect of ImmTAC version in the table at the right. * p<0.05, ** p<0.01, *** p<0.001, **** p<0.0001.

### ImmTACs overcome but don’t suppress CTL cytotoxicity

Suppressed CTL can be re-isolated from tumor cell spheroids as spheroid-infiltrating lymphocytes (SIL) to allow further investigation of their properties. To corroborate and extend the in-spheroid ImmTAC killing data (Fig. 3), we incubated 1G4 CTL with Mel624 tumor cell spheroids to induce suppressed 1G4 SIL, re-isolated them from the spheroids and determined the ability of ImmTACs to activate these suppressed cells (Fig. 4A). In the absence of ImmTAC, i.e. in response to direct 1G4 TCR engagement by NY-ESO-1_157-165_/HLA-A*0201, 1G4 CTL, i.e. the control cells that had not been incubated with spheroids, killed Mel624 target cells. However, 1G4 SIL could not, whether exogenous NY-ESO-1 peptide was added or not (Fig. 4B), corroborating the suppressed state of the 1G4 SIL. Conversely, 100pM E28 triggered cytolysis of Mel624 cells by 1G4 SIL in the presence of exogenous NY-ESO-1 peptide that was comparable to control 1G4 CTL that had not been incubated with spheroids (Fig. 4B). Even when relying on the limiting endogenous NY-ESO-1 antigen processing, 100 pM E28 induced some Mel624 target cell killing by 1G4 SIL; 1 nM E28 elicited cytolysis comparable to that of control 1G4 CTL (Fig. 4B). Consistent with the in-spheroid killing data, E28 ImmTAC thus could trigger tumor target cell killing in suppressed CTL, at a low level in response to endogenous antigen, efficiently at higher amounts of antigen or ImmTAC.

**Fig. 4.**
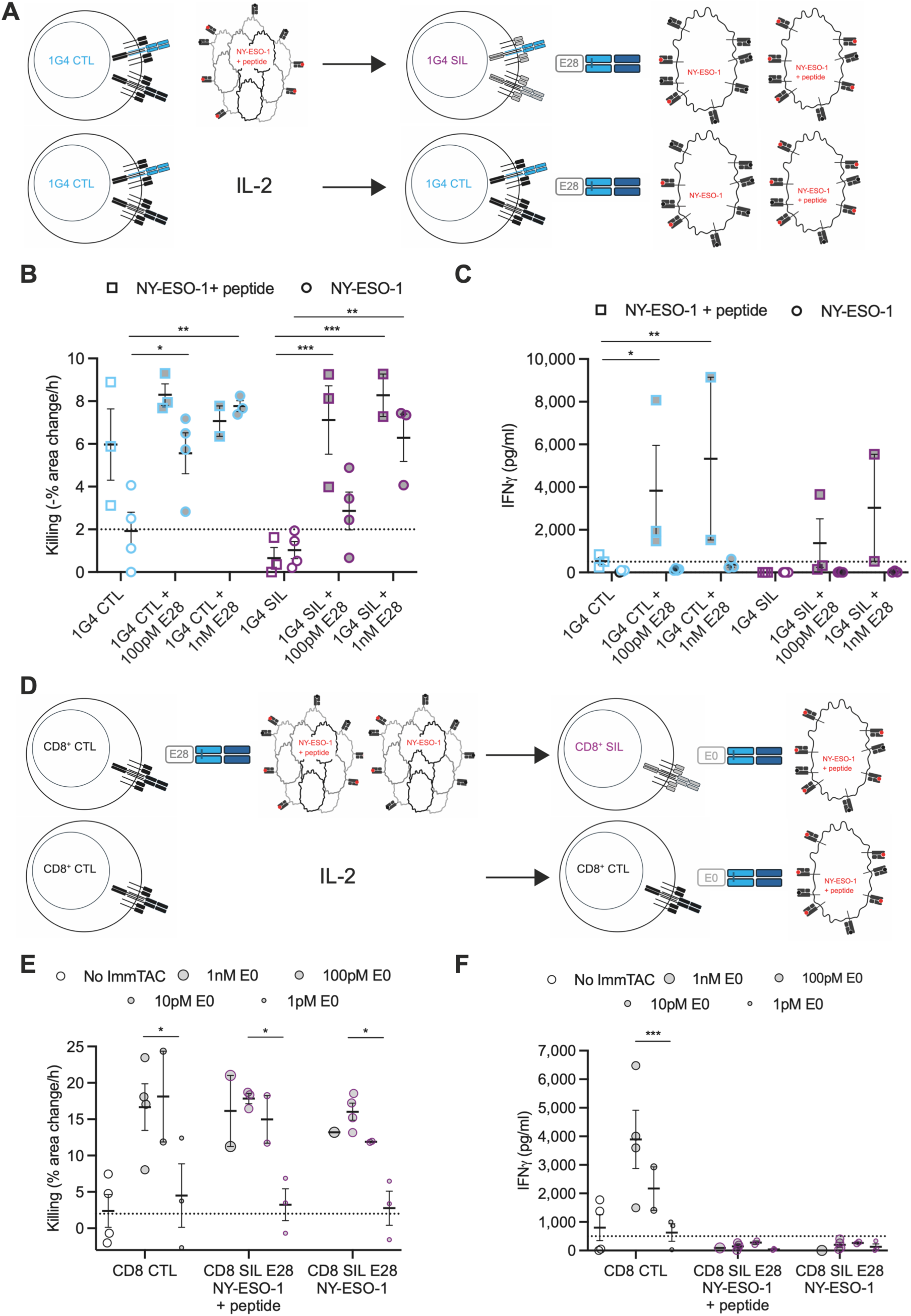
ImmTACs can redirect suppressed CTL to kill targets but do not rescue IFNγ secretion. **A** Graphical depiction of the induction of 1G4 CTL suppression and the subsequent experiment to assess activation of suppressed 1G4 SIL by ImmTAC (top row) and of the corresponding control 1G4 CTL experiment (bottom row). **B, C** Killing (B) of Mel624 tumor target cells incubated with 2 µg/ml NY-ESO-1 peptide or not by 1G4 CTL or 1G4 SIL in the absence or presence of the indicated concentration of the E28 ImmTAC and (C) IFNγ amounts in the corresponding supernatants; mean ± SEM. 4-6 independent experiments. Statistical significance determined by paired Mixed-effects analysis. **D** Graphical depiction of the experiment to test induction of suppression by ImmTAC-mediated CTL activation in spheroids and the subsequent experiment to test for such suppression (top row) and of the corresponding non-suppressed control experiment (bottom row). **E, F** Killing (E) of Mel624 tumor target cells incubated with 2 µg/ml NY-ESO-1 peptide by non-transduced CTL, previously incubated or not in spheroids incubated with 2 µg/ml NY-ESO-1 peptide or not, in the presence of 100 pM E28 ImmTAC in the absence or presence of the indicated concentration of the E0 ImmTAC and (F) IFNγ amounts in the corresponding supernatants; mean ± SEM. 1-4 independent experiments. Statistical significance determined by paired Mixed-effects analysis. * p<0.05, ** p<0.01, *** p<0.001.

However, E28 was less effective at inducing IFNγ secretion by suppressed CTL. In response to endogenous antigen processing, even at a 1 nM concentration, E28 did not induce substantial IFNγ secretion by 1G4 CTL or 1G4 SIL (Fig. 4C). In one of two experimental repeats and at saturating amounts of exogenous NY-ESO-1 peptide, E28 triggered some IFNγ secretion, albeit less than in control 1G4 CTL (Fig. 4C). Thus, at the physiologically most relevant low antigen and ImmTAC concentrations, the E28 ImmTAC could not overcome suppression of IFNγ secretion in CTL.

Antigen engagement by 1G4 CTL in spheroids overnight triggers suppression [19]. We, therefore, wanted to investigate whether comparable ImmTAC engagement in spheroids also does so. We incubated CD8^+^ CTL with Mel624 spheroids in the presence or absence of exogenous NY-ESO-1 peptide and 100 pM E28 and re-isolated the CTL as ‘CD8^+^ SIL’. We then compared activation of CD8^+^ SIL to CD8^+^ CTL that had not been incubated with spheroids in the presence or absence of exogenous NY-ESO-1 peptide, and increasing concentrations of E0 (from 1 pM to 1 nM) (Fig. 4D). E0 ImmTAC elicited CD8^+^ SIL killing of targets comparable to control CD8^+^ CTL (Fig. 4E), establishing absence of induction of suppression of cytolysis. However, the overnight incubation of 1G4 CTL with ImmTAC in spheroids prevented the CTL from subsequently secreting IFNγ, even in the presence of exogenous peptide (Fig. 4F). Thus, ImmTAC could not only rescue cytolysis in suppressed CTL (Fig. 4B) but did so without inducing a suppressed state during an overnight incubation (Fig. 4E). In contrast, such incubation induced suppression of IFNγ secretion.

To verify that TCR engagement by antigen in the spheroids is required for the induction of suppression, we incubated CD8^+^ CTL with Mel624 spheroids. Because of the diverse TCR repertoire of the CD8^+^ CTL, antigen-specific TCR engagement will be minimal. To prevent CTL reversion to a resting phenotype, spheroids were incubated with IL-2 (Fig. S4A). CD8^+^ SIL from the spheroid incubation in the absence of antigen killed Mel624 target cells in response to 100 pM E0 and secreted IFNγ as well as control CD8^+^ CTL (Fig. S4B, C), establishing absence of suppression. The induction of CTL suppression in spheroids thus required TCR engagement.

To more broadly address the ability of ImmTACs to overcome suppression, we investigated an alternate means of inducing CTL suppression, co-culture of tumor target cells with fibroblasts (Fig. S4D) to mimic cancer associated fibroblasts [27]. When Mel624 cells were co-cultured with MRC-5 fibroblasts for 16 h before the killing assay, 100 pM E28 ImmTAC no longer elicited CD8^+^ CTL killing of Mel624 cells presenting endogenous levels of NY-ESO-1 peptide (Fig. S4E). MRC-5 fibroblasts were not killed in the same experiments despite low level endogenous expression of NY-ESO-1 (not shown and Fig. S4F), removing fibroblast death as a potential confounder of the experiment. IFNγ secretion under these conditions was too low to allow a meaningful analysis. These results suggest that in a complex tumor microenvironment with a dense fibroblast network, ImmTAC may not be sufficient to rescue CTL suppression and redirect T cells to kill target tumor cells.

### ImmTACs effectively enhance the subcellular polarization of CTL interacting with tumor target cells

The execution of target cell killing by CTL requires the formation and effective maintenance of CTL target cell couples [15, 17, 19]. We therefore investigated whether ImmTACs can support such subcellular polarization of both CTL and SIL at different levels of antigen. We determined six elements of CTL tumor cell couples, i. the formation of a tight cell couple upon initial CTL tumor cell contact as a measure of CTL-perceived initial stimulus strength (Fig. S5A), ii. the time of formation of the first off-interface lamella, iii. frequency and time of translocation of the CTL over the tumor cell surface (Fig. S5B) and iv. ‘detachment’, the complete repolarization away from the tumor cell (Fig. S5C), as measures of inefficient maintenance of CTL tumor cell couples. v. F-actin clearance at the interface center (Fig. S5A) is associated with efficient delivery of lytic granules and vi. F-actin accumulation at the entire interface (Fig. S5A-C) is required for interface formation and maintenance. Incubation with ImmTACs consistently led to more stable cell couples in the form of enhanced cell couple formation (Fig. 5A), reduced translocations and detachments (Fig. 5B, C), delayed formation of the first off-interface lamellae and delayed translocations (Fig. 5D, S5D) and more effective clearance of F-actin from the interface center (Figs. 5E-H, S5E-H). Only overall interface F-actin accumulation didn’t differ (Fig. S6). E28 was consistently more effective than E0. Effects on 1G4 SILs were often more pronounced than the effects on 1G4 CTL. These polarization data closely match the ability of ImmTACs to induce effector function (Figs. 1-3). High frequencies of translocation and detachment are associated with an inability of T cells to kill [19]. While in response to saturating amounts of exogenous antigen 100 pM E28 could reverse elevated frequencies of translocation and detachment to levels observed in CTL activation, in response to endogenous antigen 100 pM E28 reduced translocation frequencies only slightly, without reaching levels seen in CTL. These data are consistent with the limited ability of E28 to induce cytolysis in SIL in response to endogenous antigen (Figs. 3D, 4B). Together these data establish that ImmTACs, similar to direct TCR engagement by MHC/peptide, support a key cellular mechanism of target cell lysis, the effective subcellular polarization of CTL as required for the release of lytic granules.

**Fig. 5.**
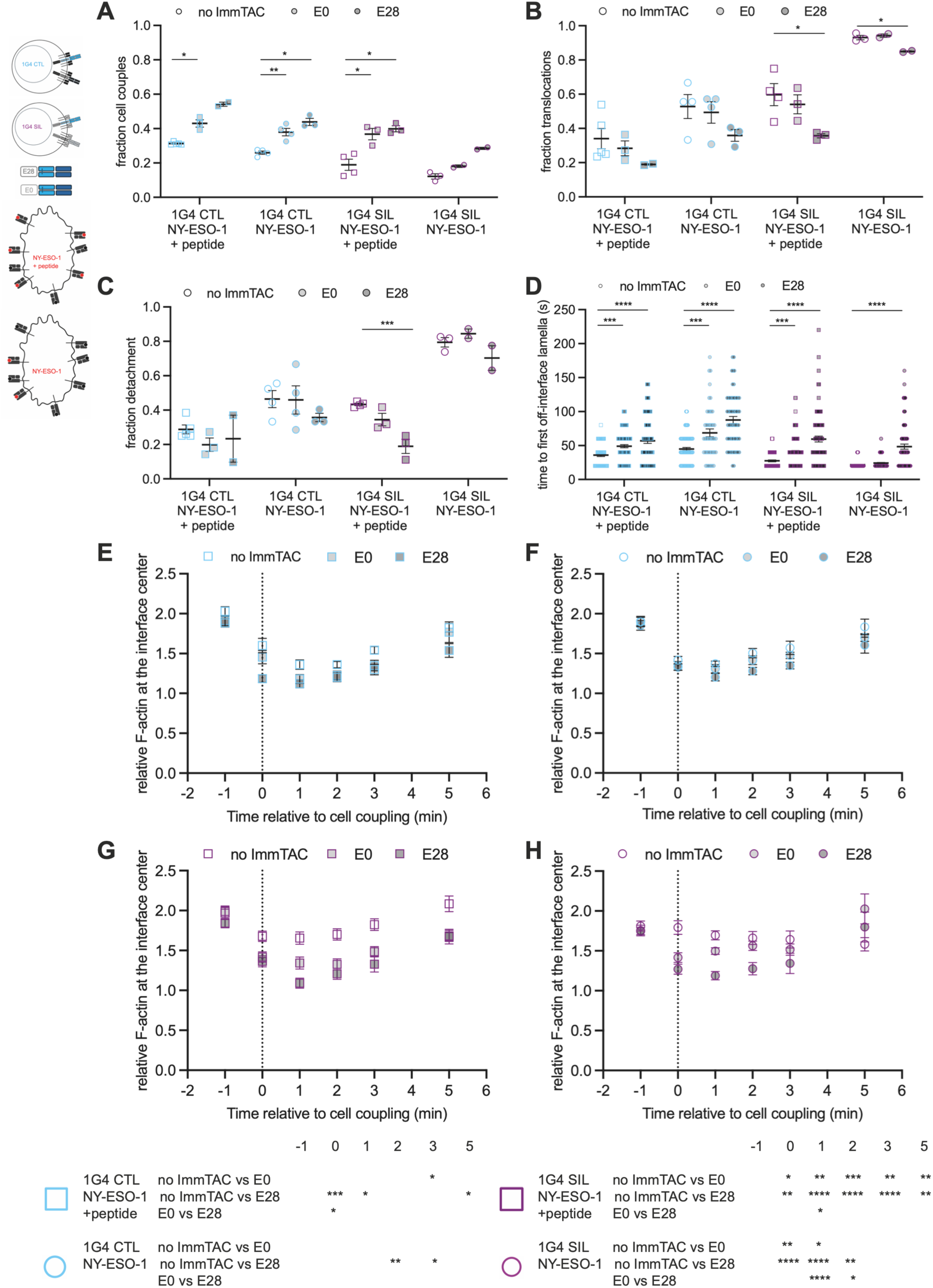
ImmTACs effectively induce cytoskeletal polarization of CTL interacting with tumor target cells. **A-D** Characterization of cell morphology in the interaction of 1G4 CTL or SIL with Mel624 cells in the presence or absence of 2 µg/ml NY-ESO-1 agonist peptide and 100 pM of the indicated ImmTAC; mean ± SEM. (A) Fraction of 1G4 CTL or SIL converting a target cell contact into a tight cell couple. Each symbol is an imaging run. (B) Fraction of CTL with a translocation. Each symbol is an imaging run. (C) Fraction of CTL with detachment. Each symbol is an imaging run. (D) Time from tight cell couple formation to first off-interface lamella. Each symbol is a cell couple. 2-5 independent experiments. Statistical significance determined by One-way ANOVA (A-C) and Kruskall-Wallis test (D). **E-H** F-actin accumulation at the center of the interface between 1G4 CTL (E, F) and 1G4 SIL (G, H) expressing F-tractin-GFP and Mel624 cells in the presence (E, G) or absence (F, H) of NY-ESO-1 agonist peptide and 100 pM of the indicated ImmTAC relative to F-actin in the entire cell and to the time of tight cell coupling; mean ± SEM. Pooled data from 2-3 independent experiments. Single cell data in Fig. S5E-H. Statistical significance determined by Two-way ANOVA and given in the table at the bottom right. * p<0.05, ** p<0.01, *** p<0.001, **** p<0.0001.

### ImmTACs trigger effective calcium signaling but no short-term changes in coregulatory receptor expression

The elevation of the cytoplasmic calcium concentration is a key proximal signaling event in CTL activation that closely correlates with IFNγ secretion [18]. Using the calcium-sensitive dye Fura-2 we determined calcium signaling in 1G4 CTL, CD8^+^ CTL and 1G4 SIL in response to 100 pM E28 as compared to direct engagement of the 1G4 TCR by MHC/peptide (Fig. 6A). Measurable calcium signaling in 1G4 CTL required the incubation of Mel624 cells with exogenous NY-ESO-1 peptide. 100 pM E28 further enhanced calcium signaling triggered by direct engagement of the 1G4 TCR with MHC/peptide (Fig. 6B, S7A). As expected, no calcium signaling was observed in CD8^+^ CTL in the absence of the E28 ImmTAC (Fig. 6C, S7B). Additionally, calcium signaling triggered by Mel624 cells in the presence of exogenous NY-ESO-1 peptide and 100 pM E28 was smaller in CD8^+^ CTL than in 1G4 CTL (Fig. 6B, C). These data suggest that the E28 ImmTAC is more effective in triggering calcium signaling when acting in parallel to direct TCR engagement by MHC/peptide rather than in isolation. Such enhanced signaling did not translate into enhanced effector function (Fig. S1C, D) as further discussed below. In 1G4 SIL, no calcium signaling was detected in the absence of E28, whether the Mel624 cells were incubated with exogenous NY-ESO-1 peptide or not, consistent with the suppressed state of the 1G4 SIL (Fig. 6D, S7C). E28 could trigger effective calcium signaling in the presence of exogenous NY-ESO-1 peptide, consistent with its ability to overcome 1G4 SIL suppression under these conditions (Fig. 4B).

**Fig. 6.**
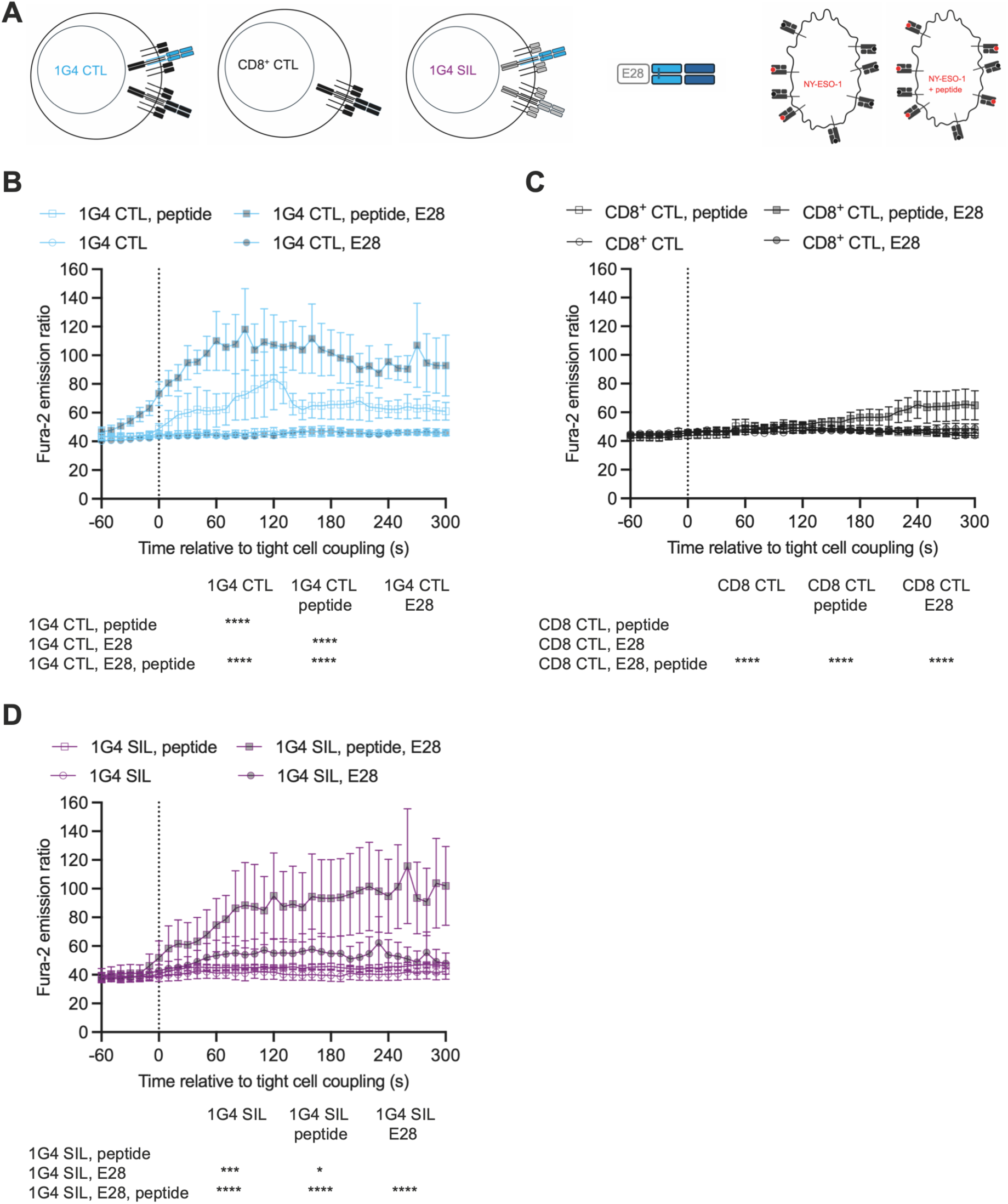
E28 ImmTAC triggers the elevation of the CTL cytoplasmic calcium concentration more effectively than direct engagement of a TCR by MHC/peptide. **A** CTL and target cells used in B-D. **B-D** Ratio of the Fura-2 emission upon excitation at 340 nm over 380 nm of 1G4 CTL (B), CD8^+^ CTL (C) or 1G4 SIL (D) activated by Mel624 cells in the presence or absence of 2 µg/ml NY-ESO-1 agonist peptide and 100 pM E28; mean of independent experiments ± SEM. 3-4 independent experiments. Statistical significance determined by paired Mixed-effects analysis and given in the tables at the bottom. Single cell data are given in Fig. S7. * p<0.05, *** p<0.001, **** p<0.0001.

To determine whether ImmTAC treatment on the overnight time scale of our experiments changes protein expression, we determined the expression of 17 T cell markers focused on coregulatory receptors in the interaction of CD8^+^ CTL with Mel624 cells incubated with exogenous NY-ESO-1 peptide in the presence of 100 pM E28 for 4 h and 16 h. No substantial changes occurred, neither in the percentage of CD8^+^ CTL positive for a given marker, nor in the mean fluorescence intensity (Fig. 7, S8A-E). The lack of substantial changes upon ImmTAC treatment is confirmed in a cluster analysis (Fig. S8F-H). A small fraction of the CTL displayed a more exhausted phenotype characterized by higher expression of inhibitory receptors and lower expression of TCF1 and SLAMF6. However, this population was restricted to strongly stimulated CTL, CTL in the presence of IL-2 or ImmTAC for 4 h. On the time scale of the execution of CTL effector function investigated here, effects of ImmTACs thus were predominantly posttranslational.

**Fig. 7.**
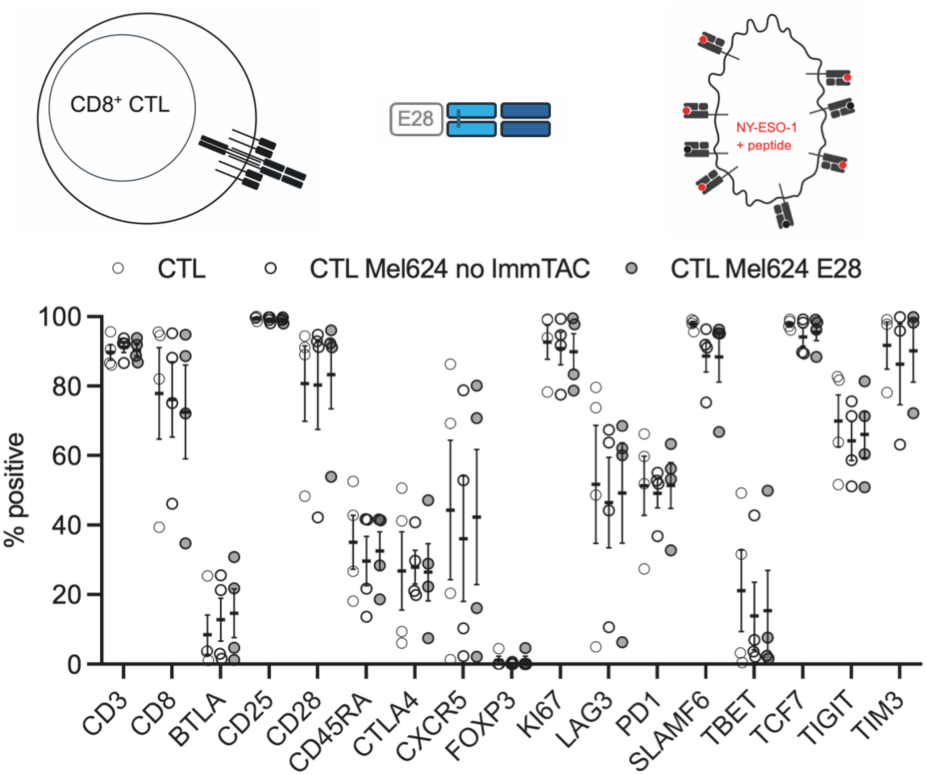
E28 ImmTAC does not trigger substantial changes in T cell marker expression on the time scale of the execution of CTL effector function. Percentage of CD8^+^ CTL positive for the indicated marker in CD8^+^ CTL, CD8^+^ CTL after 4 h of incubation with Mel624 cells in the presence of exogenous NY-ESO-1 agonist peptide with or without 100 pm E28. 4 independent experiments. Representative flow cytometry data, mean fluorescence intensity data and data after 16 h of incubation with Mel624 cells in Fig. S8.

## Discussion

### ImmTACs as tunable T cell engagers

ImmTACs use an anti-CD3ε scFv antibody fragment to trigger TCR signaling. In the use of agonist antibodies, varying the antibody affinity commonly results in a bell-shaped dose response curve where both low and high affinities trigger suboptimal cellular effector function [25, 28]. In this paper, we performed experiments using ImmTAC molecules engineered with a series of anti-CD3ε scFv antibody fragments of varying affinity. We observed that, in response to NY-ESO-1 peptide pulsed targets, a similar bell-shaped dose response curve exists for ImmTACs, with the most potent induction of IFNγ secretion at an anti-CD3ε affinity of 35 nM [12]. (Fig. 1C). However, when we reduced the stimulus strength to more closely mimic conditions expected in vivo, by relying on endogenous NY-ESO-1 antigen processing and using fully active (Fig. 1D, E) or suppressed (Fig. 3D) CTL as effectors, the dose response curve shifted. Under these antigen-limiting conditions, T cell activation was most effective using the highest affinity anti-CD3ε scFV antibody fragment, E42. A similar shift in the dose response curve was found in vitro when reducing the affinity of the TCR moiety of the ImmTAC for MHC I/peptide [12]. Adjusting the anti-CD3ε affinity of an ImmTAC thus may be an effective strategy to adapt the ImmTAC to the physiological circumstances of therapeutic application. Potential losses in specificity with increasing affinity [12] need to be considered in doing so. These considerations should also apply to the design of new TCR engagers [20] based on synthetic MHC/peptide binders [20–22] which currently have a comparatively low affinity for the MHC/peptide complex (in the low nM range) that may require compensation by a higher affinity anti-CD3ε scFV antibody fragment.

### ImmTAC function under conditions of CTL suppression

In therapeutic settings, ImmTACs must function under conditions of CTL suppression in the tumor microenvironment. We have used tumor cell spheroids here to reflect such suppression. In the presence of a high amount of antigen, both ImmTACs with an intermediate and a high affinity for CD3ε (E28, E42) effectively elicited killing of tumor target cells in spheroids (Fig. 3B) and upon isolation of suppressed T cells from spheroids (Fig. 4B). Using the high affinity E42 ImmTAC some cytolytic activity was even observed against Mel624 cells presenting low levels of endogenous NY-ESO-1 peptide. Antigen engagement under suppressive conditions triggers CTL exhaustion. ImmTACs were not only able to overcome CTL suppression in a spheroid system (Fig. 3B, D, 4B), they did so without inducing suppression of the cytolytic ability of CTL (Fig. 4E). However, ImmTACs could not overcome suppression of IFNγ secretion, nor suppression of cytolysis in the context of stromal cells (Fig. S4C, E). Out data suggest that in vivo translation may benefit from patient stratification by antigen amount and CAF numbers or from combination therapies.

In the extrapolation of in vitro data to ImmTAC function in a therapeutic setting, other CD8^+^ T cell effector functions and CD4^+^ T cells also need to be considered. In contrast to the ImmTAC-dependent preservation of cytolytic function, the ability to secrete IFNγ was readily lost in a suppressive environment (Fig. 4C, F). It thus seems unlikely that tumor-resident CTL from patients treated with ImmTACs make much IFNγ. Our work here has not considered the effect of ImmTACs on CD4^+^ T cells. In vitro ImmTACs can trigger target cell killing and cytokine secretion by CD4^+^ T cells [29]. The effect on regulatory T cells has not been determined either, but previous work showed that in vitro ImmTAC-triggered CD8^+^ T cell function is insensitive to the presence of regulatory T cells [29]. For a more comprehensive extrapolation of mechanisms of ImmTAC function in vivo, future work will need to focus on CD4^+^ T cells, helper and regulatory, under suppressive conditions and on CD8^+^ and CD4^+^ T cells in conjunction.

### The mechanism of CTL activation by ImmTACs is similar to that of direct TCR engagement by MHC/peptide

Activating a T cell by replacing the engagement of its TCR by MHC/peptide complexes with synthetic approaches can lead to substantial alterations in T cell signaling. For example, proximal T cell signaling triggered by chimeric antigen receptors is substantially less efficient [30]. The total size of a receptor ligand complex at the interface between a T cell and an activating antigen presenting cell is important for signaling efficiency [31]. This could be relevant here, as the total size of a MHC/peptide-ImmTAC-CD3ε complex might be larger than that of a MHC/peptide-TCRα/β complex. When assayed as the ability to trigger cytolysis and IFNγ secretion, T cell activation by ImmTACs at physiologically realistic concentrations displayed a comparable peptide sensitivity and extent of effector function than T cell activation by direct TCR engagement with MHC/peptide (Fig. 2). Moreover, ImmTACs triggered the same effective execution of a series of CTL subcellular polarization steps required for efficient cytolysis as direct TCR engagement (Fig. 5). ImmTACs thus closely mimic physiological T cell activation. There was one notable difference. Under the limiting T cell activation conditions investigated here, the elevation of the intracellular calcium concentration triggered by direct TCR engagement was inefficient (Fig. 6B, D). Calcium signaling triggered by ImmTAC in CTL not expressing a tumor antigen-reactive TCR was equally inefficient (Fig. 6C). However, calcium signaling triggered by ImmTAC in 1G4 CTL was substantially elevated, in particular in SIL (Fig. 6B, D). Such elevation, however, did not correlate with more efficient CTL effector function (Fig. S1C, D). These data suggest that TCR engagement at a single point of the TCR complex, be it the α/β dimer or CD3ε, is equally efficient. However, engagement of the complex at two points, α/β dimer plus CD3ε be it on the same TCR molecule or different TCR molecules at the same cellular interface, can trigger more efficient proximal signaling. We speculate that comparable engagement of coregulatory receptors in these two scenarios prevents the more efficient TCR engagement from triggering stronger effector function. The molecular mechanism of these signaling differences needs to be resolved in the future. However, the strong similarity in the sensitivity and cellular mechanism of T cell activation by ImmTACs in comparison to direct TCR engagement suggests that mechanisms of the physiological regulation of T cell activation, for example control by coregulatory receptors, may apply to T cell function in patients treated with ImmTACs. ImmTACs thus may benefit from combination with therapies that have already proven effective in the activation of endogenous T cells, such as checkpoint blockade.

### Materials and Methods Human blood samples

Blood buffy coats from anonymous donors were purchased from NHS-BT with human work approved by the London-Riverside Research Ethics Committee under reference number 20/PR/0763.

### Antibodies

Antibodies and staining reagents used are described in the order: antigen, fluorescent label, clone, supplier, dilution/concentration, RRID:

Panel for flow cytometry
Human CD25, BV421BD, M-A251, BD Bioscience, 1:100, RRID:AB_11154578
Human CD25, BV421BD, 2A3, BD Bioscience, 1:100, RRID:AB_2738555
Human CD8α, Pacific Blue, RPA-T8, BioLegend, 1:25, RRID:AB_493111
Human CD28, BV480, CD28.2, BD Bioscience, 1:100, RRID:AB_2739512
Human CD4, Starbright V570, RPA-T4, Bio-Rad, 1:50, RRID:AB_3099780
Human CD3ε, Super Bright 645, OKT3, eBioscience, 1:20, RRID:AB_2662368
Human CD185 (CXCR5), BV785, J252D4, BioLegend, 1:50, RRID:AB_2629527
Human CD366 (TIM-3), Super Bright 600, F38-2E2, eBioscience, 1:100, RRID:AB_26882087
Human CD45RA, R718, HI100, BD Bioscience, 1:100, RRID:AB_2916420
Human CD279 (PD-1), APC-Cy7, EH12.2H7, BioLegend, 1:100, RRID:AB_10900982
Human CD45, NovaBlue 610-30S, 2D1, eBioscience, 1:100, RRID:AB_3098021
Human CD352 (SLAMF6/Ly108), biotin, REA339, Miltenyi, 1:30, RRID:AB_2657676
Streptavidin, Star Bright Blue 675, Bio-Rad, 1:85
Human CD272 (BTLA), BB700, J168-540, BD Bioscience, 1:50, RRID:AB_2743523
Human CD152 (CTLA-4), PE-Fire 640, BNI3, BioLegend 1:20, RRID:AB_2924561
Human TIGIT, PE-vio770, REA1004, Miltenyi, 1:100, RRID:AB_2751339
Human CD223 (LAG-3), PE-Fire 810, 7H2C65, BioLegend, 1:100, RRID:AB_2927867
Human CD14, RB545, M5E2, BD Bioscience, 1:100, RRID:AB_2691229
Human CD19, RB545, SJ25C1, BD Bioscience, 1:100, RRID:AB_2688586
Human Ki-67, BV711, Ki-67, BioLegend, 1:100, RRID:AB_11218996
Human Foxp3, PE, 259D, BioLegend, 1:50, RRID:AB_492983
Human Foxp3, PE, PCH101, eBioscience, 1:50, RRID:AB_1518782
Human TCF1, Alexa Fluor 647, 7F11A10, BioLegend, 1:50, RRID:AB_2566619
Human T-bet, RB780, O4-46, BD Bioscience, 1:100, RRID:AB_2688938
Live Dead Red, Invitrogen, 1:20,000

### Proteins

ImmTAC proteins with varying anti anti-CD3ε scFv effector domains were expressed in the BL21 (DE3) Rosetta pLysS strain, refolded from inclusion bodies and purified as previously described [12, 32]. Purity was checked by reducing and non-reducing SDS-PAGE and concentrations were assessed by A280 measurement and extinction coefficients derived from sequence by the inbuilt DNAdynamo algorithm.

### Human Cell culture

Human cell culture was executed as previously described [19] with minor adjustments. Human A375 melanoma (RRID:CVCL_0132) and Mel624 melanoma (RRID:CVCL_8054) cells transfected to express mCherry or tdTomato [19, 33] were maintained in high glucose DMEM or RPMI-1640, respectively, with 10% FBS, 2 mM Glutamine, 1 mM pyruvate (DMEM/RPMI complete medium) supplemented with 250 μg/ml Hygromycin. MRC-5 human embryonic lung fibroblasts (RRID:CVCL_0440) were transfected to express GFP and maintained in DMEM complete medium.

Blood buffy coats were obtained from healthy donors. PBMC were isolated by density gradient centrifugation using Ficoll-Paque^TM^ (Sigma-Aldrich). PMBC were cryopreserved at a concentration of 3×10^7^ cells/ml. To isolate CD8^+^ T cells, buffy coat cryopreserved vials were thawed, the cells were washed twice with RPMI 1640 with 10% FBS, 2mM L-glutamine, 50 µM β-mercaptoethanol and resuspended in ice cold MACS buffer. CD8^+^ T cells were purified by magnetic enrichment for CD8^+^ cells using CD8 MicroBeads (Miltenyi Biotech). CD8^+^ T cells were activated using CD3/CD28 Dynabeads (Life Technologies) at bead-to-cell ratio of 1:1 in human IL-2 medium (X-VIVO 15, serum-free hematopoietic cell medium, with 2 mM L-Glutamine and gentamicin (Lonza) supplemented with 5% Human AB serum (Valley Medical), 10 mM neutralized N-acetyl L-Cysteine (Sigma-Aldrich), 50 μM β-Mercaptoethanol (Gibco, ThermoFisher), and 30 U/ml rh-IL-2 (NIH/NCI BRB Preclinical Repository – human IL-2 medium) and incubated overnight at 37°C and 6% CO2.

The 1G4 TCR, stabilized with an additional disulfide bridge in the constant domains [26], was expressed in primary human T cells using a pHR_SFFV -based lentiviral vector (RRID_Addgene79121) with an expression cassette of alpha chain-P2A-beta chain-P2A-GFP (as a sorting marker) or F-tractin-GFP (for F-actin imaging)[19]. For the generation of lentiviral particles, HEK 293T cells (RRID:CVCL_0063; Lenti-X 293T cells, Takara) were maintained in DMEM complete medium. 1.5×10^6^ Lenti-X 293T cells were seeded in 5 ml DMEM complete medium in 60×15 mm Primaria culture plates (Corning) 24 h before transfection. Cells were transfected with a total of 4.5 µg plasmid DNA using Fugene HD (Promega): 0.25 µg envelope vector pMD2.G (RRID:Addgene_12259), 2 µg of packaging plasmid pCMV-dR8.91 (Creative Biogene), and 2.25 µg of the pHR_SFFV-based transfer vector. 48 h after transfection, virus containing medium was collected and filtered through a 0.45 µm nylon filter. For lentiviral infection of CTL, after 24 h of setting up the primary human T cell culture, 1×10^6^ T cells were mixed with 500-700 µl lentivirus-containing medium in a 24-well plate medium in presence of 8 µg/ml Polybrene (Sigma-Aldrich) and centrifuged for 1.5 h at 2500rpm, 32°C. After spinduction primary CD8^+^ T cells were resuspended in human IL-2 medium. Cells were maintained at densities of less than 2×10^6^ cells/ml and if necessary, spilt back to a density of 0.5-1×10^6^ cells/ml.

### Spheroids and SIL

Experiments with spheroids and SIL were executed as previously described [19] with minor adjustments. Mel624 tdTomato cells were resuspended at a concentration of 1×10^5^ cells/ml, mixed with Matrigel (Corning) at 4°C, seeded in a 24-well plate at a final concentration of 500 cells per Matrigel dome, and left to solidify for 10 min at 37°C. 2 ml RPMI complete medium was added to each well and cells incubated at 37°C for 11 days.

For the generation of spheroid-infiltrating lymphocytes (SIL), each Matrigel dome was washed twice in PBS and incubated for 1 h with 1 ml of Cell Recovery Solution (Corning) at 4°C. Spheroids were collected in a 15 ml Falcon tube and pulsed with NY-ESO-1 peptide at a final concentration of 2 μg/ml or with 100 pM of the E28 ImmTAC for 1 h each or left unpulsed. Spheroids were re-embedded in Matrigel together with 5×10^6^ 1G4 or non-transduced CD8^+^ CTL per Matrigel dome. Matrigel domes were dissolved for analysis of spheroid-infiltrating T cells after 16 h: Spheroids were washed twice in PBS and incubated with 1 ml of Cell Recovery Solution (Corning). Spheroids were collected, washed through a 40 μm sieve and then disaggregated to retrieve T cells in 1 ml of 10% FBS in PBS with 1 mM CaCl_2_ and 0.5 mM MgCl_2_ for immediate FACS sorting.

To determine CTL infiltration into spheroids by live cell imaging, spheroids were dissociated from Matrigel and resuspended into fresh Matrigel at a concentration of ∼8 spheroids/µl. 50 µl of the spheroid-Matrigel suspension was separated into Eppendorf tubes, followed by the addition of 500,000 CTL per tube. 50 µl of Matrigel, containing spheroids and T cells, was plated into each well of a 24-well black, glass base, sterile Sensoplate^TM^. After Matrigel had set, 1 ml of Fluorobrite medium (ThermoFisher) was added to each well, containing 1.5 µM DRAQ7 viability dye. Images were acquired every 2 h post-plating CTL with spheroids in 3 µm z steps from the bottom of the spheroid to its widest point, usually 40 steps, for 16 h using a Leica SP8 AOBS confocal microscope with a 10x HC PL Fluotar lens (NA=0.3).

Spheroids and SIL were segmented using a custom Fiji Image J script as described before [34]. Distance of each T cell from the spheroid surface was automatically calculated using the script.

### Cytolysis and IFNγ secretion

Cytolysis and IFNγ secretion were determined as previously described [19] with minor adjustments. For imaging-based cytotoxicity assays, the IncuCyte™ Live Cell analysis system and IncuCyte™ ZOOM software (Essen Bioscience) were used to quantify target cell death. 1×10^6^ Mel624 or A375 cells transfected to express the fluorescent protein mCherry or tdTomato were either untreated or pulsed for 1 h with 2 µg/ml of HLA-A*02:01 NY-ESO-1_157-165_ (SLLMWITQC), HLA-A*02:01 MART-1_26-35_ with the A27L mutation (‘ELA’)(ELAGIGILTV) or the indicated concentration of an ImmTAC. Cells were suspended in 5 ml Fluorobrite medium (ThermoFisher) with 10% FBS, 2 mM L-glutamine, 50 µM 2-mercaptoethanol to a concentration of 10,000 cells/50 μl. MRC5 cells were transfected to express GFP. Cells were plated in each well of a 384 well Perkin-Elmer plastic-bottomed plate and incubated for 4 h to adhere. For Mel624 MRC5 co-cultures cells were seeded at a 1:1 ratio and incubated for 16 h. 40,000 CTL, non-transduced or that had been FACS sorted for GFP, were added per well to the plate in 50 μl Fluorobrite medium, yielding a 4:1 effector to target ratio. Images were taken every 15 min for 14 h at 1600 ms exposure using a 10x (NA=0.3) lens. The total red object (mCherry/tdTomato target cell) area (µm^2^/well) was quantified at each time point. To determine MRC5 killing green object area was quantified. A change in red/green area was determined as the linear gradient of the red/green object data at its steepest part for 6 h. The CTL killing rate was calculated as the difference in such change in red/green area in the presence and absence (to account for tumor cell proliferation) of CTL in the same 6 h time window.

To determine IFNγ secretion, supernatants at the end of the imaging-based cytolysis assays were collected and frozen. The human IFNγ OptEIA ELISA Kits (BD Biosciences) was used according to manufacturer’s instructions. Briefly, wells of a 96-well Maxisorb Nunc-Immuno plate (ThermoFisher) were coated with 100 µl anti-human IFNγ monoclonal antibody diluted in 0.1 M Na_2_CO_3_ coating buffer at a 1:250 dilution and incubated overnight at 4°C. Plates were blocked using PBS with 10% FBS for 1h. Samples were diluted in ELISA dilution buffer at 1:3 to 1:10 and incubated for 1h at room temperature. 100 µl of working detectors (Biotinylated anti-human IFNγ monoclonal antibody + Streptavidin-horseradish peroxidase conjugate) were incubated for 1 h at room temperature before detection. Absorbance at 450 nm and 570 nm was measured within 20 min. Samples were measured in duplicate or triplicate.

### Imaging of CTL and SIL

Imaging of CTL and SIL was executed as previously described [19]. Prior to imaging, CTL and SIL were resuspended in ‘imaging buffer’ (10% FBS in PBS with 1 mM CaCl_2_ and 0.5 mM MgCl_2_). As target cells, 1×10^6^ Mel624 melanoma cells were pulsed with the indicated peptide (as listed above) at a final concentration of 2 µg/mL for 1 h or left unpulsed. Glass bottomed, 384-well optical imaging plates (Brooks life science systems) were used for all imaging experiments. Imaging of F-actin distributions and CTL morphology was done at 37°C using an Olympus IXplore SpinSR confocal system incorporating a Yokogawa CSU-W1 SoRa spinning disk. A 60x oil-immersion lens (NA=1.5) was used for all imaging experiments, unless otherwise stated. Images were acquired for 15 min. Every 20 s, a z-stack of 53 GFP images (0.25 µm z-spacing) was acquired, as well as a single, mid-plane differential interference contrast (DIC) reference image.

For imaging the elevation of the cytoplasmic calcium concentration, 1G4 or non-transduced CD8^+^ T cells were incubated with 2 µM Fura-2 AM (Molecular Probes) for 30 min at room temperature in imaging buffer. CTL were washed twice thereafter. Because of limiting cell numbers 1G4 SIL were only washed once. T cells were activated with Mel624 APCs as described above and three images were acquired every 10 s for 15 min, one bright field image, one fluorescence image with excitation at 340 nm and one fluorescence image with excitation at 380 nm. Imaging data were acquired at 37°C using a 40x oil objective (NA=1.25) on a Leica DM IRBE-based wide filed system equipped with Sutter DG5 illumination and a Photometrics Coolsnap HQ2 camera.

### Analysis of live cell imaging data

Using Fiji/ImageJ [35, 36] for analysis of DIC images, tight cell couple formation was defined as the first time point at which a maximally spread immune synapse formed, or 40 s after initial cell contact, whichever occurred first. To assess CTL and SIL morphology in cell couples with tumor target cells, every DIC frame after tight cell couple formation was assessed for the presence of off-synapse lamellae, defined as transient membrane protrusions pointing away from the immune synapse, followed by retraction. To determine CTL translocation over the tumor cell surface, the position of the immune synapse on the tumor target cell was compared to the position at cell coupling. If the T cell had migrated by a distance greater than the diameter of the immune synapse, this was classed as translocation. The formation of a uropod such that it is bound to the tumor cell at the interface rather than pointing away from it together with lamellae at the CTL side opposite of the interface was classed as ‘detachment’.

For analysis of F-actin distributions as imaged with F-tractin-GFP, enrichment at the interface and interface center were measured relative to F-actin across the entire cell using Fiji/ImageJ. F-tractin-GFP enrichment across the interface relative to the entire cell was determined in the maximum intensity z-projection of the GFP z-stack. The interface was defined as the 10% of the area of the T cell closest to the T cell/target cell interface. The average fluorescence intensity of the interface area and the entire cell were measured at four to six timepoints after subtracting the mean of ten background fluorescence readings. F-tractin-GFP enrichment at the centre of the interface relative to the entire cell was determined in the midplane of the GFP z-stack of the T cell of interest. The area of the interface centre was defined as the middle third of the interface as defined above, that is as 3.5% of the total cell midplane area. The average fluorescence intensity of the interface center and the entire cell were measured at four to six timepoints after subtracting the mean of ten background fluorescence readings.

For analysis of the Fura-2 imaging experiments using Fiji/ImageJ, rolling ball background fluorescence was subtracted from the fluorescence data and the ratio of the Fura-2 images upon excitation at 340 nm versus 380 nm was calculated and multiplied by 100 to fit into the 8-bit display scale. Average ratio within a circular region of interest of the dimensions of the T cell was determined over time for each T cell.

### Flow cytometry

Flow cytometry was executed as previously described [19]. 21-color panel: 1×10^6^ cells were first stained with a 1:10,000 dilution of the LIVE/DEAD™ Fixable Red Dead Cell Stain (ThermoFisher) for 15-30 min in the dark at room temperature and washed with FACS buffer (PBS, 0.5% BSA, 2.5 mM EDTA). Fc receptors were blocked with 10 µl of a 1:20 dilution of Fc Blocker (ThermoFisher) in the dark at room temperature for 10 min. 40 µl of biotinylated antibody was added and incubated at 4°C for 30 min. The sample was washed with FACS buffer and incubated with 10 µl of a 1:20 dilution of Monocyte Blocker (BioLegend) in the dark at room temperature for 10 min. 40 µl of an antibody master mix against cell surface proteins as listed in the antibodies section was added for the total volume of 50 µl, incubated at 4°C for 30 min and washed with FACS buffer. Cells were fixed and permeabilized using the True-Nuclear™ Transcription Factor Buffer Set (BioLegend). Cells were stained for intracellular proteins with an antibody master mix as listed in the antibodies section for 30 min at 4°C. Cells were washed with permeabilization buffer and resuspended in FACS buffer and kept at 4°C until the next day. For flow cytometry data acquisition, samples were run on a four-laser Cytek® Aurora system (4L V/B/YG/R) on low to medium flow rates (15 - 30 µl/min) and analyzed using the SpectroFlo® software with autofluorescence extraction and live unmix during sample acquisition. The data for the unmixed samples were processed using the FlowJo software (v.10.10.0). Positive gates were set using fluorescence-minus-one data.

For cluster analysis, samples were gated for live, singlet, CD45^+^CD3^+^CD19^−^CD14^−^CD8^+^ cells. The FlowJo 10.10.0 clean plugin was used to reduce technical noise. Files for experimental repeats were concatenated with the FlowJo concatenation tool. CD8^+^ cells for each experimental repeat were down sampled to 20,000 cell and analyzed for CD25, CD28, CD185, CD366, CD45RA, CD279, CD352, CD272, CD152, TIGIT, CD223, Ki-67, FOXP3, TCF1, T-BET. t-SNE (t-Distribution Stochastic Neighbor Embedding) was employed to map the high-dimensional cellular data into a two-dimensional space using: iteration: 1000, perplexity: 30, K-nearest neighbors’ algorithm: Exact (vantage point tree), and gradient algorithm: Barnes-Hut. Cells were clustered by implementing FlowSOM v. 3.0.18 with 10 metaclusters and 10 × 10 grid size, all other parameters were left at the default settings.

## Acknowledgements

We acknowledge support from the University of Bristol Flow Cytometry and Wolfson BioImaging core facilities.

## Funding sources

This work was supported by grants from the MRC (MR/W006308/1 to TG for the GW4 BIOMED MRC DTP), the Wellcome Trust (338308/Z/25/Z to AH-W for the Dynamic Molecular Cell Biology DTP) and Immunocore (to MS, CJH and CW). AbA and AmA were supported by the Ministry of Education of Saudi Arabia.

## Author contributions

LH, AbA, AmA Investigation, Formal Analysis, Methodology, Writing Review & Editing; TG, AH-W Investigation, Formal Analysis; JC, GP, AC Investigation; MS, CJH, Formal Analysis, Funding Acquisition, Writing Review & Editing, CW Investigation, Formal Analysis, Visualization, Funding Acquisition, Writing Initial Draft, Writing Review & Editing

## Conflict of Interest

MS and CJH are full-time employees and shareholders at Immunocore.

## Notes

### Summary of Updates

Missing references to previous work from our laboratory have been added to the methods section. The manuscript is otherwise unchanged from the earlier version.

