## Supplementary figures for "ImmTACs overcome cytotoxic T cell suppression"

1 School of Cellular and Molecular Medicine, University of Bristol, Bristol, BS8 1TD, UK
2 Department of Life Sciences, University of Bath, Bath, BA2 7AY, UK
3 ImmunoCore Ltd., Abingdon, OX14 4RY, UK
4 Newcastle Drug Discovery Group, Translational and Clinical Research Institute, Newcastle University, Newcastle upon Tyne, NE2 4HH, UK
### equal contribution

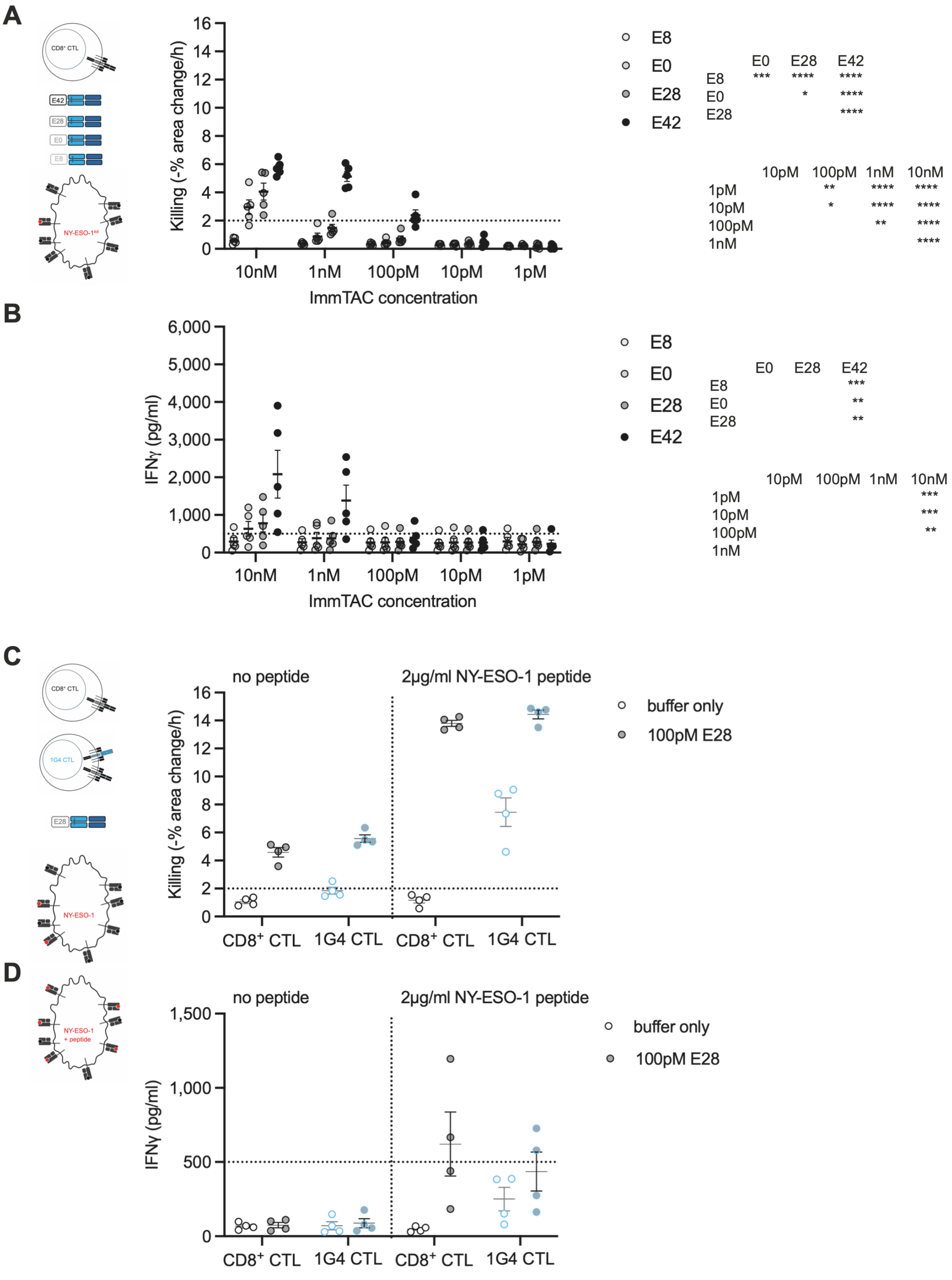


**Fig. S1 ImmTAC-dependent CTL effector function is reduced upon knockdown of NY-ESO-1**

**A, B** Killing (A) of Mel624 NY-ESO-1 knockdown tumor target cells by non-transduced CTL in the presence of the indicated concentration of the given ImmTAC version and (B) IFNγ amounts in corresponding supernatants; mean ± SEM. 5 independent experiments.

Statistical significance determined by paired Two-way ANOVA and given as main effects of ImmTAC version and concentration in the two tables at the right. **C, D** Killing (C) of Mel624 tumor target cells in the presence of exogenous NY-ESO-1 agonist peptide or not by non-transduced $CD8^+$ or 1G4 CTL in the presence or absence of 100 pM of the E28 ImmTAC and (D) IFN$\gamma$ amounts in corresponding supernatants; mean ± SEM. 4 independent experiments.

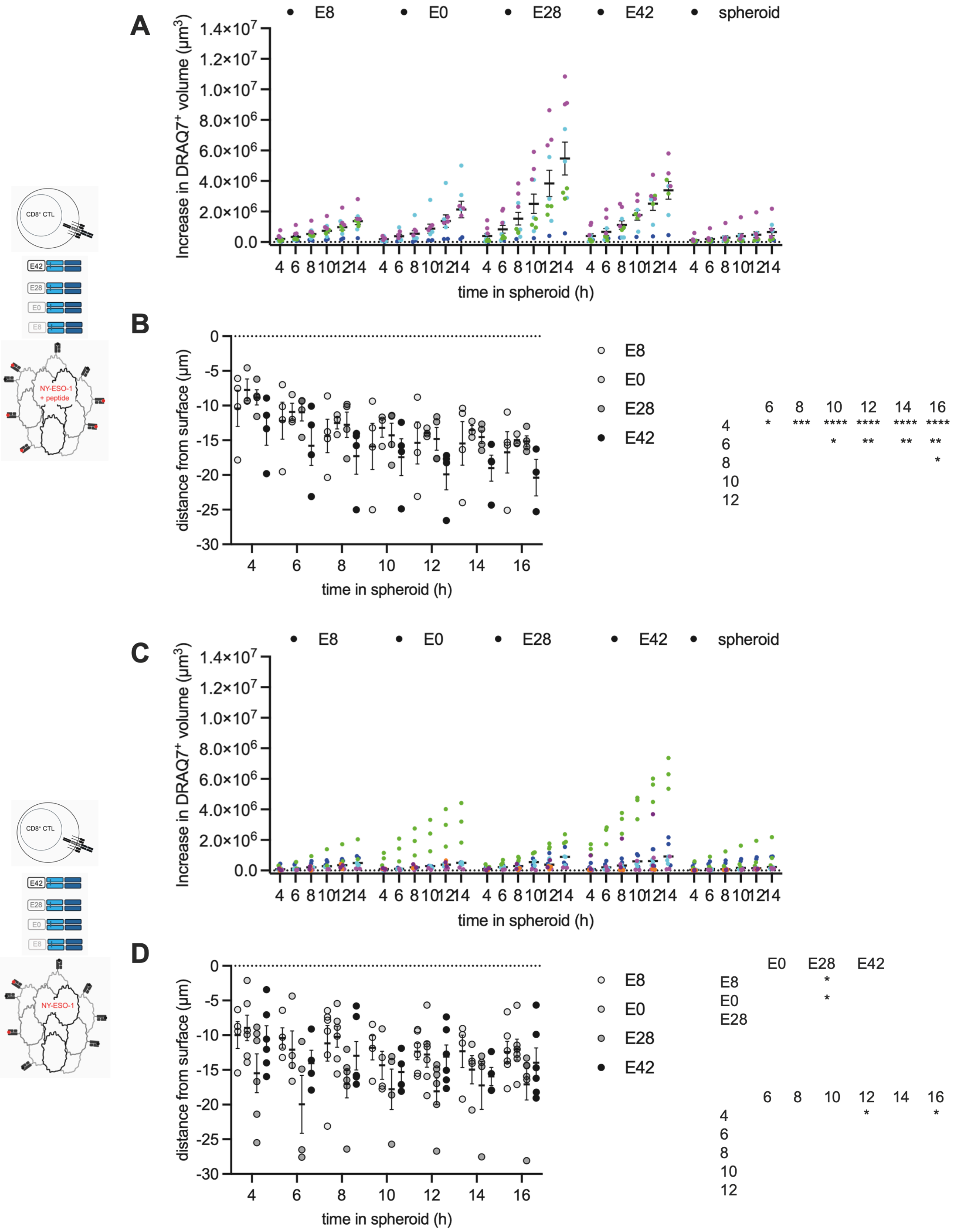


**Fig. S2 ImmTACs mediate target cell killing in tumor cell spheroids**
**A** Single spheroid data for Fig. 3B. Spheroids from the same experiment share their color across all experimental conditions. **B** SIL infiltration depths for the same experiments as in Fig. 3B; mean ± SEM. Statistical significance determined by paired Mixed-effects analysis and given as main effect of time in the table at the right. **C** Single spheroid data for Fig. 3D as in A. **D** SIL infiltration depths for the same experiments as in Fig. 3D; mean ± SEM. Statistical significance determined by paired Mixed-effects analysis and given as main effects of ImmTAC version and time in the tables at the right.

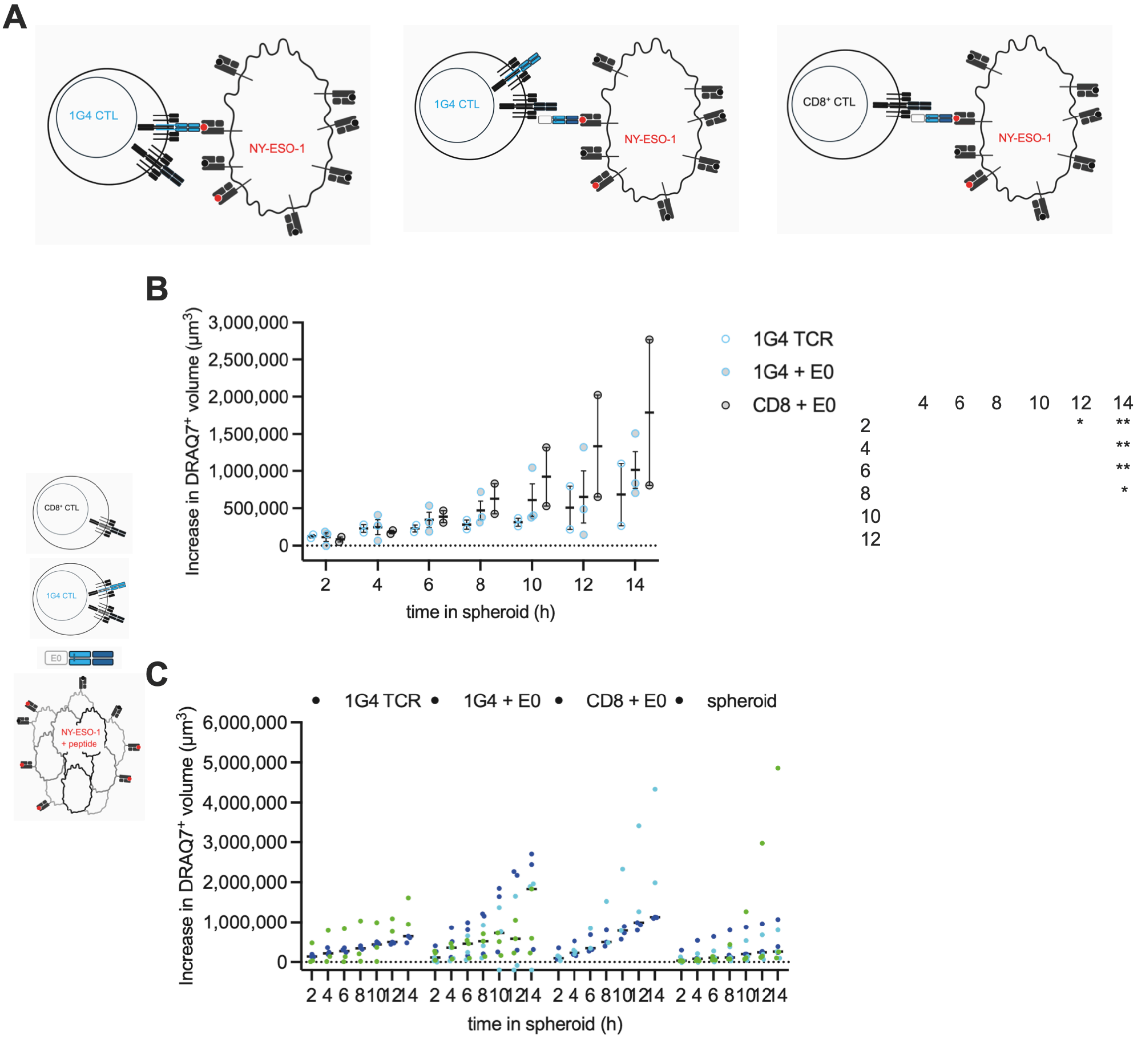


| | 4 | 6 | 8 | 10 | 12 | 14 |
|---|---|---|---|---|---|---|
| 2 | | | | | * | ** |
| 4 | | | | | | ** |
| 6 | | | | | | ** |
| 8 | | | | | | * |
| 10 | | | | | | |
| 12 | | | | | | |

**Fig. S3 Comparable target cell killing efficiency of tumor cell spheroids elicited by CTL activated via MHC/peptide or ImmTACs**

**A** Graphical depiction of the direct activation of 1G4 CTL by NY-ESO-1$_{157\text{-}165}$/HLA-A*0201 versus the indirect activation of 1G4 or non-transduced CTL via ImmTACs recognizing the same NY-ESO-1$_{157\text{-}165}$/HLA-A*0201 complex. **B** 1G4 and non-transduced CTL cocultured with Mel624 spheroids incubated with 2 µg/ml NY-ESO-1 peptide with and without 100 pM E0 ImmTAC. Each data point is an independent experiment (N=2, 3) with a total of 6-9 spheroids analyzed per condition. Statistical significance determined by paired Mixed-effects analysis and given as main effect of time in the table at the right. Single spheroid data are given in C. **C** Single spheroid data for B. Spheroids from the same experiment share their color across all experimental conditions. * $p<0.05$, ** $p<0.01$.

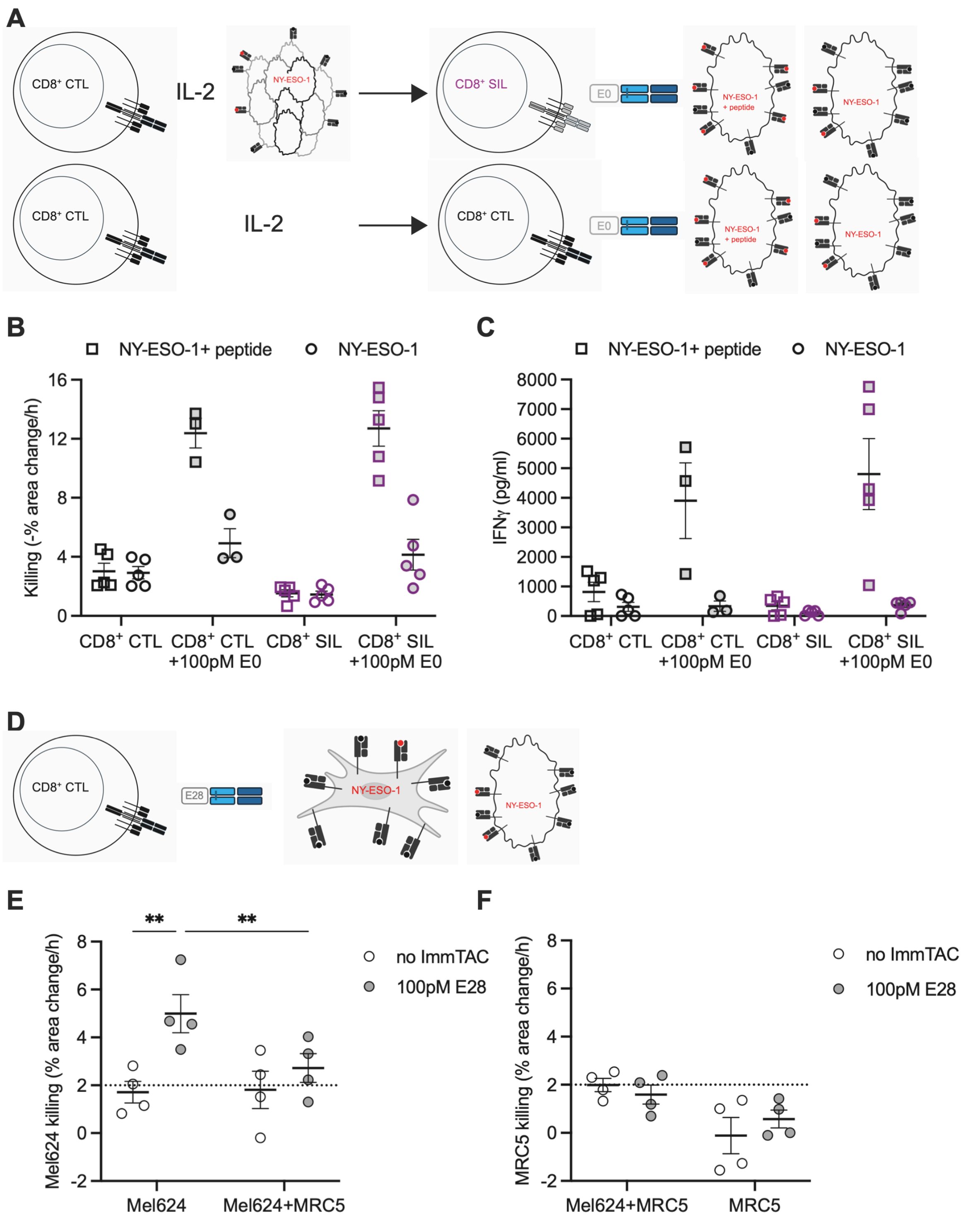


**Fig. S4 TCR engagement with antigen is required for the induction of CTL suppression in spheroids**

**A** Graphical depiction of the experiment to test induction of suppression by CTL incubation in spheroids in the absence of antigen and the subsequent experiment to test for such suppression (top row) and of the corresponding non-suppressed control experiment (bottom row). **B, C** Killing (B) of Mel624 tumor target cells with or without incubation with 2 µg/ml NY-ESO-1 peptide by non-transduced CTL previously incubated or not in spheroids without antigen exposure in the absence or presence of 100 pM E0 ImmTAC and (C) IFNγ amounts in the corresponding supernatants; mean ± SEM. 3-5 independent experiments. **D** Graphical depiction of the experiment to test suppression of CTL function in response to ImmTAC by CAFs. **E, F** Killing of Mel624 tumor target cells (E) or MRC5 CAF (F) with or without incubation with 100 pM E28 by non-transduced CTL. A negative killing rate indicates faster cell proliferation than control; mean ± SEM. 4 independent experiments. Statistical significance determined by paired Two-way ANOVA. ** $p<0.01$.

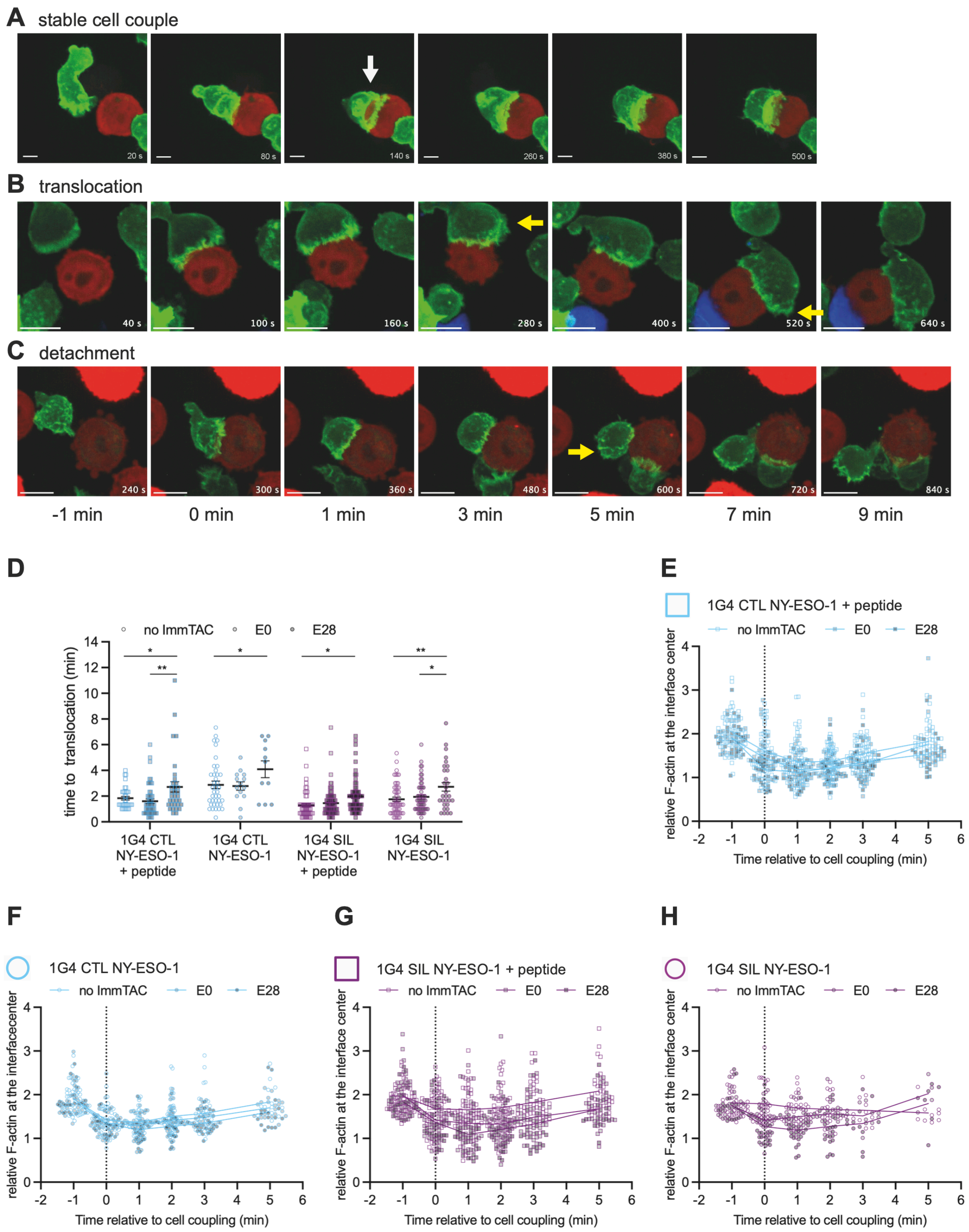


**Fig. S5 ImmTACs effectively induce cytoskeletal polarization of CTL interacting with tumor target cells**

**A-C** Maximum projections of 3D imaging data for the representative phenotypes as indicated of the interaction of 1G4 CTL transduced to express F-tractin-GFP (green) and $CD8^+$ CTL (blue) with Mel624 target cells (red) at the indicated times relative to the formation of a tight cell couple. The white arrow denotes F-actin clearance at the interface center, the yellow arrow off interface lamellae. Scale bar=5μm. **D** Time to translocation in the interaction of 1G4 CTL or SIL with Mel624 cells in the presence or absence of 2 μg/ml NY-ESO-1 agonist peptide and 100 pM of the indicated ImmTAC; mean ± SEM. Part of the same experiments as Fig. 5 A-D. Statistical significance determined by One-way ANOVA. **E-H** Single cell data for Fig. 5E-H. On average 36 (26-64) cell couples analyzed per condition. * $p<0.05$, ** $p<0.01$

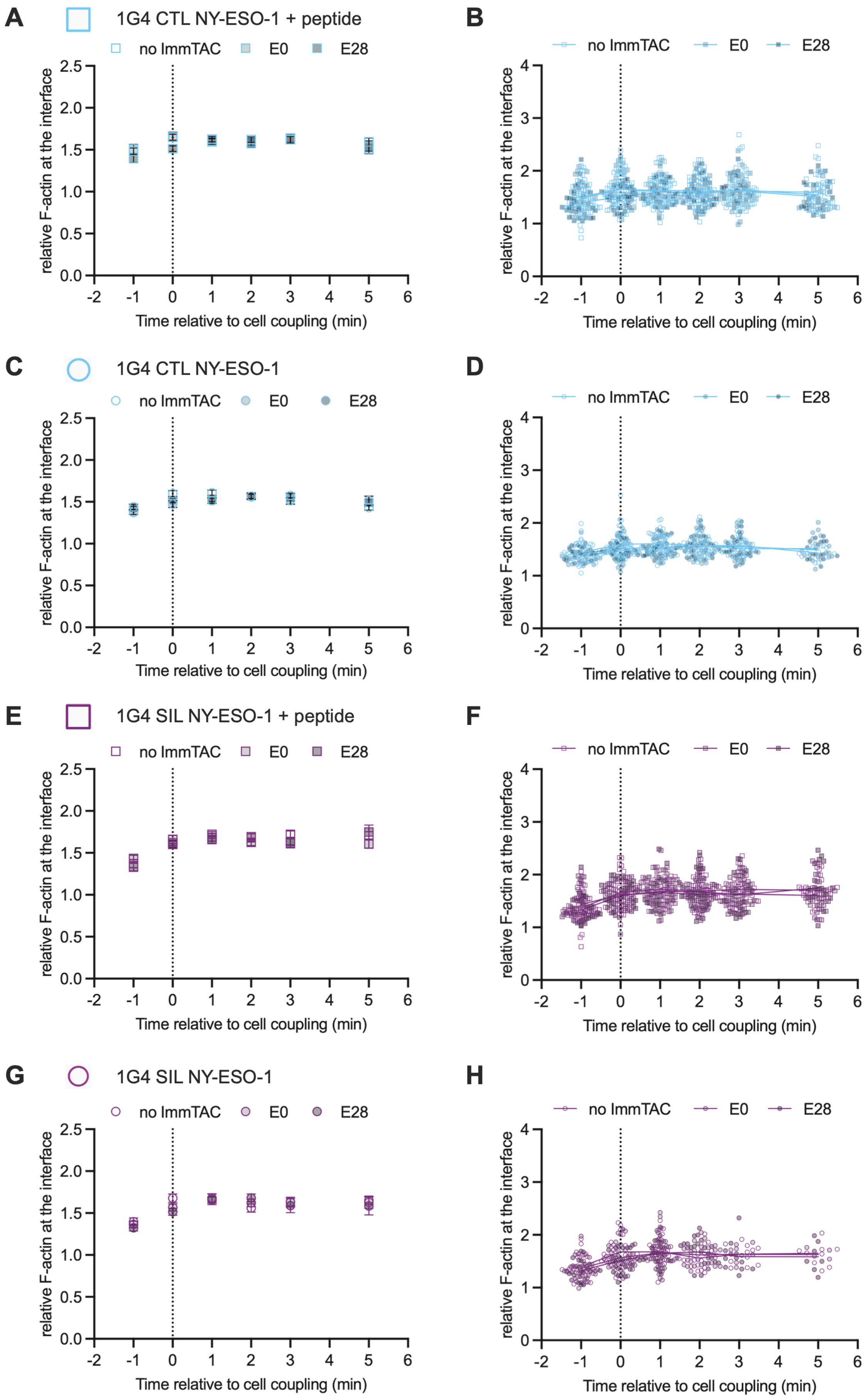
A
1G4 CTL NY-ESO-1 + peptide
no ImmTAC
E0
E28
relative F-actin at the interface
Time relative to cell coupling (min)
B
C
1G4 CTL NY-ESO-1
D
E
1G4 SIL NY-ESO-1 + peptide
F
G
1G4 SIL NY-ESO-1
H

**Fig. S6 ImmTACs effectively induce cytoskeletal polarization of CTL interacting with tumor target cells**

**A-H** F-actin accumulation at the entire interface between 1G4 CTL (A-D) and 1G4 SIL (E-H) expressing F-tractin-GFP and Mel624 cells in the presence (A, B, E, F) or absence (C, D, G, H) of NY-ESO-1 agonist peptide and 100 pM of the indicated ImmTAC relative to F-actin in the entire cell and to the time of tight cell coupling. Pooled data from 2-3 independent experiments. A, C, E, G Experiment averages; mean ± SEM. B, D, F, H Corresponding single cell data. On average 36 (26-64) cell couples analyzed per condition.

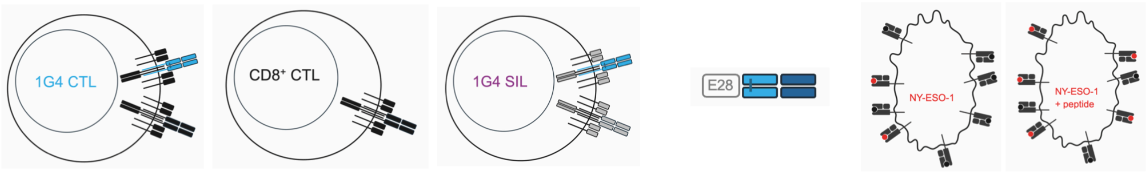


**A**

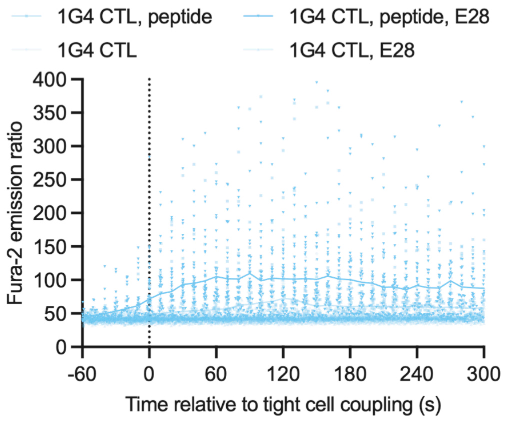


| | 1G4 CTL | 1G4 CTL peptide | 1G4 CTL E28 |
|---|---|---|---|
| 1G4 CTL, peptide | **** | | |
| 1G4 CTL, E28 | | **** | |
| 1G4 CTL, E28, peptide | **** | **** | **** |

**B**

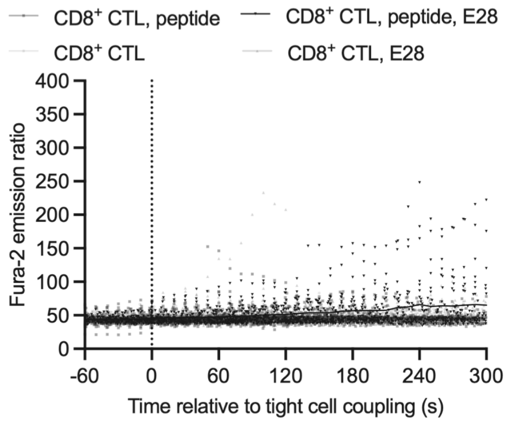


| | CD8 CTL | CD8 CTL peptide | CD8 CTL E28 |
|---|---|---|---|
| CD8 CTL, peptide | | | |
| CD8 CTL, E28 | | * | |
| CD8 CTL, E28, peptide | **** | **** | **** |

**C**

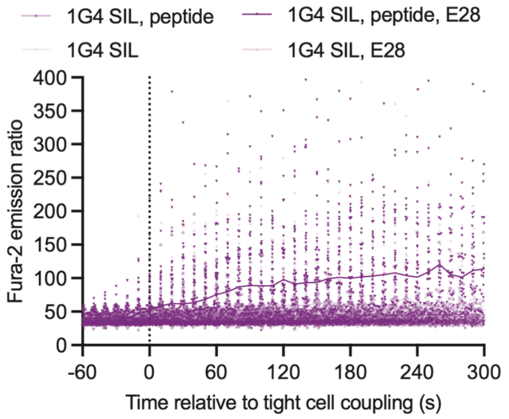


| | 1G4 SIL | 1G4 SIL peptide | 1G4 SIL E28 |
|---|---|---|---|
| 1G4 SIL, peptide | | | |
| 1G4 SIL, E28 | **** | **** | |
| 1G4 SIL, E28, peptide | **** | **** | **** |

**Fig. S7 E28 ImmTAC triggers the elevation of the CTL cytoplasmic calcium concentration more effectively than direct engagement of a TCR by MHC/peptide**
**A-C** Single cell data for Fig. 6B-D, respectively, as pooled over all independent experiments of a given condition. 32-117 cells analyzed per condition. Statistical significance determined by paired Mixed-effects analysis and given in the table at the bottom. * $p<0.05$, **** $p<0.0001$.

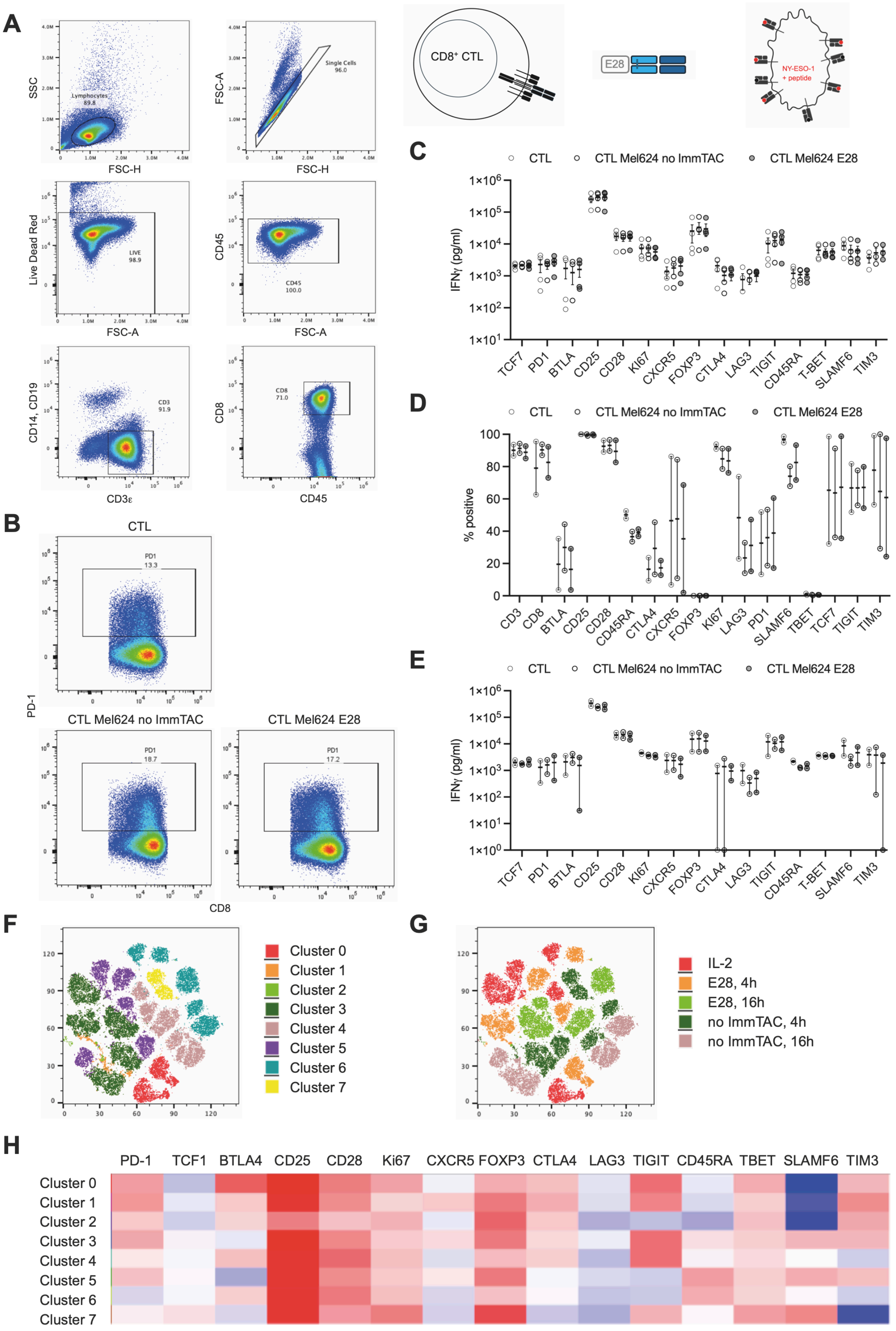


**Fig. S8 E28 ImmTAC does not trigger substantial changes in T cell marker expression on the time scale of the execution of CTL effector function**

**A** Gating strategy **B** Representative flow cytometry data. **C** Mean fluorescence intensity data for Fig. 7. **D, E** Percentage (D) of $CD8^+$ CTL positive for the indicated marker and (E) mean fluorescence intensity in $CD8^+$ CTL, $CD8^+$ CTL after 16 h of incubation with Mel624 cells in the presence of exogenous NY-ESO-1 agonist peptide with or without 100 pm E28. 2 independent experiments. **F-H** Cluster analysis of the same data as in Fig. 7 as, (F) a t-SNE blot of clusters identified, (G) overlay of the five experimental conditions and (H) expression of the indicated markers.
